# Next-Generation Soybean Haplotype Map as A Genomic Resource for Enhanced Trait Discovery and Functional Analysis

**DOI:** 10.64898/2026.03.24.713798

**Authors:** Aamir W. Khan, Dadakhalandar Doddamani, Qijian Song, Tri D. Vuong, Sushil S. Chhapekar, Heng Ye, Vanika Garg, Rajeev K. Varshney, Henry T. Nguyen

## Abstract

We present a global soybean haplotype map generated from whole-genome sequencing of 1,278 *Glycine max* and *Glycine soja* accessions, comprising 11.37 million SNPs and 2.05 million short insertions and deletions. This map (GmHapMap-II) captures unprecedented worldwide genetic diversity, reflecting the broad extent of the global soybean gene pool. Population structure analyses revealed six geographically distinct subpopulations that affected the linkage and shaped the recombination. The haplotype variation map was used to identify novel genomic regions associated with crude protein content on chromosome 15 that were not detected by a lower SNP density array. LD-based haplotype analysis revealed a superior haplotype for crude protein content. The constructed haplotype map enabled detailed characterization of haplotype diversity and copy number polymorphism at the SCN-associated *rhg-1 and Rhg-4* loci, revealing both novel haplotype structures and germplasm lines with elevated CNV relative to previously characterized genotypes. We employed the HapMap matrix for a multi-class variations ML-based genomic prediction approach to predict phenotypes for SCN and catalogued the gene-centric haplotypes in a user-friendly database. The analysis revealed the extent of deleterious alleles present in the soybean germplasm and how breeders have deployed beneficial alleles and purged deleterious ones. The haplotype map will serve as a major genomic resource for trait-based mapping, enhancing efforts in the genomics-enabled development of improved cultivars.

## Introduction

Soybean [*Glycine max* (L.) Merr.] is a crop of significant economic importance as it serves as a primary source for both animal feed protein and oil content. The allopolyploid genome, sequenced for the first time in 2010, comprises approximately 1.1 billion base pairs across 20 chromosomes, with roughly 58,000 protein-coding genes (Schmutz et al., 2010). This genomic architecture, combined with the plant’s capacity for symbiotic nitrogen fixation and production of valuable proteins and oils, makes soybean an essential subject for studying plant evolution, adaptation and improving crop yields through biotechnology. Soybean production is severely hampered by biotic and abiotic stresses, and it is crucial to identify beneficial superior alleles for improved varieties (Fang et al., 2026; Tian et al., 2025). Meeting projected 2050 protein demands requires exploiting the substantial genetic diversity preserved in germplasm collections, yet this diversity remains poorly characterized at the genomic level. While previous resequencing efforts have catalogued millions of single-nucleotide polymorphisms (SNPs), reliance on a single linear reference genome has limited detection of structural variants and resulted in incomplete linkage disequilibrium, reducing statistical power in genome-wide association studies and leaving substantial trait heritability unexplained (Zhou et al., 2015; Fang et al., 2017). Future progress in soybean breeding relies on deeper insights into genetic diversity, especially at the allelic level, present in worldwide germplasm resources. Genomic resources play an integral part in this strategy, and generating comprehensive resources can lead to more informed decisions in breeding based on genomics (Varshney et al., 2021).

Recent advances in long-read sequencing technologies have enabled dramatic improvements in reference genome quality. The improved soybean reference assembly Wm82.a5, constructed using long-read technology, exhibits substantially enhanced contiguity and completeness compared to previous versions, enabling more comprehensive detection of structural variants and complex genomic rearrangements (Garg et al., 2023). However, the full potential of this improved assembly for population-scale variant discovery and trait dissection remains unrealized. The declining cost of genome sequencing, driven by advances in next-generation sequencing technologies, has opened unprecedented opportunities to comprehensively identify genetic variants across complete genomes from multiple individuals within a species. For soybean, numerous germplasm scale sequencing efforts have been accomplished, resulting in uncovering genetic relationships and identifying millions of variations in local and global germplasms. This involved a large number of individuals sequenced from the USA (Bayer et al., 2022; Valliyodan et al., 2021; Valliyodan et al., 2016), China (Lam et al., 2010; Liu et al., 2020; Zhou et al., 2015; Fang et al., 2017), Canada (Torkamaneh et al., 2021; Torkamaneh et al., 2017), Korea (Kim et al., 2021). Despite all these efforts, most genetic and association analyses are conducted using the Soy50KSNP array panel due to its ease of use and manageable size of the genotype data (Song et al., 2013). This has yielded several important and notable quantitative trait loci (QTLs) used for candidate gene discovery in seed composition and abiotic/biotic stresses (Hwang et al., 2014; Han et al., 2015; Zhang et al., 2019). However, functional gene characterization studies and gene validation require complete sequence information, which is lacking in low-density genotyping arrays. To overcome this, there is a need to develop a comprehensive haplotype variation map that combines the genetic diversity of different germplasms and provides high-resolution genetic variations to pinpoint candidate genes associated with traits. Developing a haplotype map with global genetic variations catalogued in a single panel requires collating sequence data from lines across multiple local germplasm collections. This is accomplished by deploying a homogeneous variant calling workflow that utilizes a consistent variant identification strategy and using a single reference genome.

Haplotype maps provide a systematic framework for characterizing genomic variation patterns within a species by identifying sequence variants, their allele frequencies, and the linkage disequilibrium structure between variants across populations (Chia et al., 2012). Haplotypes, which are specific genomic blocks, emerge when sequence variants are inherited together in a highly correlated fashion, a phenomenon known as linkage disequilibrium (LD). In our quest to update and expand the reference set of segregating diversity in soybean, we integrated previously reported resequencing panels with soybean lines from six previous large-scale sequencing efforts. This includes resequencing data from soybean lines, originating on six continents, associated with domestication panels and improvement, including wild relatives, landraces, breeding lines and elite cultivars. The main objective of this study was to develop a haplotype map that improves the resolution and mapping power for genome-wide association studies conducted in important traits, including the seed composition traits. We added a resequencing of an additional 142 soybean lines at high coverage, including a diverse panel of ancestral lines, milestone lines, lines associated with seed composition traits displaying varying seed protein content and high sucrose-associated lines.

Our comprehensive variant catalog revealed extensive variations in landraces and wild species lines. The haplotype map provided higher resolution for association and fine mapping uncovering more significant genomic regions associated with protein content revealing superior haplotypes for important traits. Gene-centric haplotype analysis identified one unique functional haplotype across *rhg-1* gene associated with SCN resistance. We demonstrate practical breeding applications through genomic prediction and optimal haplotype identification. We have developed a soybean variation and haplotype database providing a user-friendly interface for extracting specific haplotypes from a large set of variation catalogues. This high-resolution haplotype map provides a foundational resource for soybean genetics and breeding, enabling improved genomic selection, targeted allele mining, and accelerated genetic improvement of this critical global protein crop.

## Results

### A global soybean HapMap

To create a global haplotype map for soybean (GmHapMap-II), we analyzed 1,278 re-sequenced soybean lines. This collection comprised 1,116 previously sequenced accessions, representing national and regional core collections, together with 162 newly sequenced accessions specifically for this study. Together, these samples are considered representative of the global soybean gene pool (**Figure 1A**).

**Figure 1:**
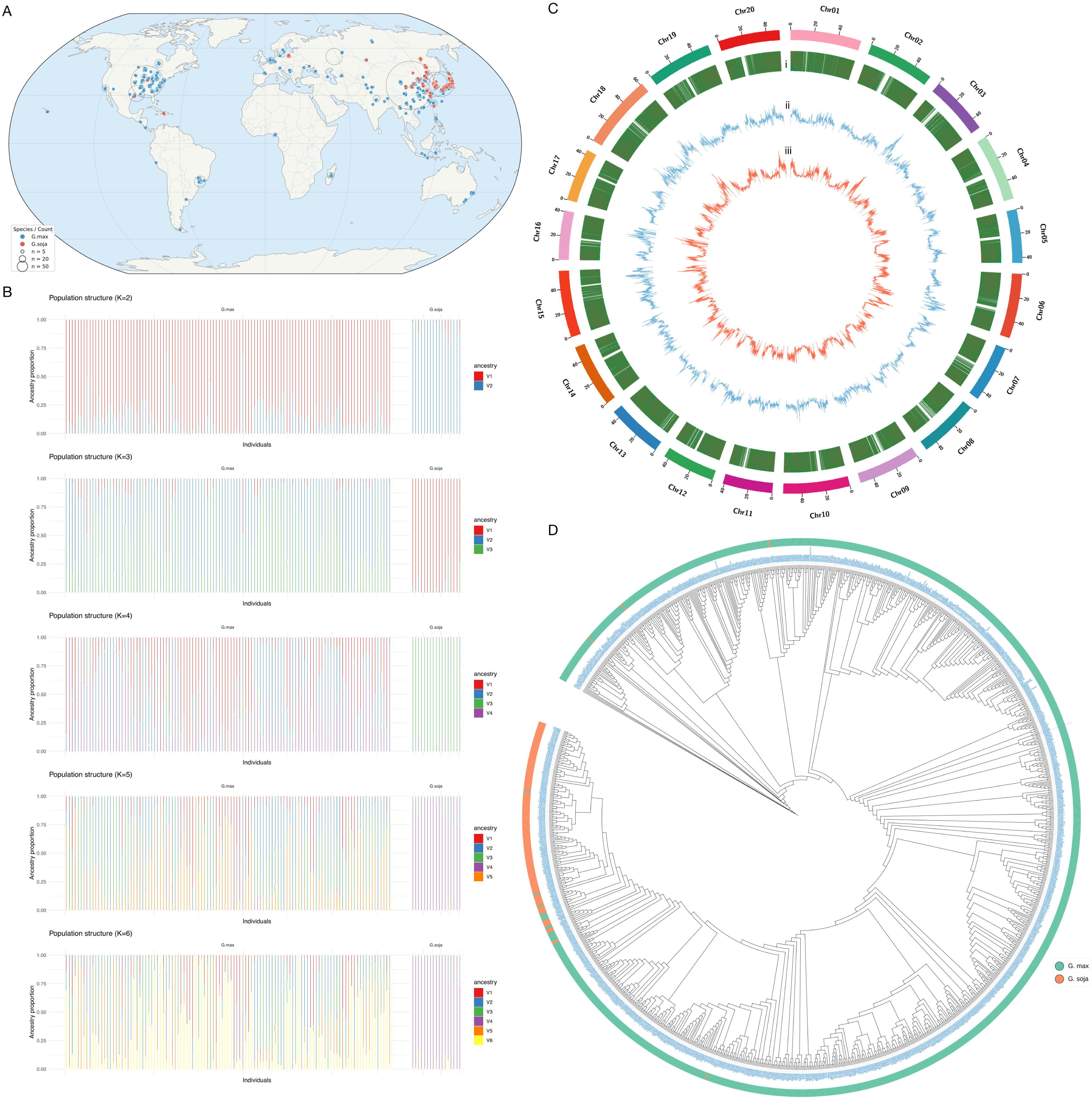
Global diversity in the Haplotype map. (A) Geographic distribution of the 1,252 *G. max and G. soja* accessions used to construct the global variation map. (B) Population structure inferred from ADMIXTURE analysis, showing subpopulation differentiation and admixture patterns across multiple k-values. (C) Circos visualization of genome-wide variation, including gene-density (i), SNP density (ii), and InDel density (iii), each calculated in 1-Mb windows across the genome. (D) Phylogenetic tree depicting evolutionary relationships among all soybean accessions based on high-quality SNPs.

The sequencing effort produced ∼206.15 billion paired-end high-quality reads (ranging from 100 to 150 base pairs), totaling ∼27.45 trillion base pairs (**Supplementary Table S1**). This provided an average sequencing depth of 19.3X. To ensure consistent variant calling, all data was processed using a unified pipeline based on GATK4. We collated whole-genome resequencing data for a total of 1,278 soybean lines, representing a global collection of soybeans from countries including the USA, China, Korea, Japan, Brazil and Australia, among the major ones (**Figure 1A; Supplementary Table S1; Supplementary Table S2**). The lines include a collection of 1,119 cultivated lines and 159 *G. soja* lines which constituted 701 landraces, 268 cultivar lines, 28 breeder lines, 141 wild-type and 140 others. Short-reads data from these accessions were mapped to the near-gapless Williams 82 v5 genome (Garg et al., 2023), which yielded an average mapping coverage of 98.9% of the genome (**Supplementary Table S2**). Analysis of the germplasm collection revealed subpopulation admixture in the US national set, which covers a larger genetic diversity (**Figure 1B**). The variant calling resulted in the identification of 50.97 million raw SNPs which were further filtered down with GATK hard filtration parameters to 11.37 million (MAF>0.01) high-quality SNPs for most downstream population analyses **(Figure 1C; Supplementary Table S3**). Over for all the soybean samples, the mean SNPs per sample was ∼1,664,955. The highest number of SNPs was observed for PI 424007 (3,822,504) and the least for PI 548631 (62,403; **Supplementary Table S4**). The annotations of these variants showed that the highest proportion of SNPs were present in the intergenic regions (9,600,721; 84.47%) followed by intronic SNPs (1,346,700; 11.85%; **Supplementary Table S5**). The missense variations were comparatively lower in frequency undermining the phenomenon of conserved sequences within the gene boundaries. The analysis revealed high effect variations such as start lost, stop gained and stop lost were present in fewer variants and affected a total of 419 genes. Further examining these high-effect variations, we identified a total of 13,841 high-effect SNPs across the soybean germplasm lines, with the highest number observed on Chr 18 (1165; **Supplementary Table S6**). Our analysis of high-effect SNPs and InDels, including loss-of-function (LOF) variants, showed that several of these variants affect important trait-associated genes, such as disease resistance genes, genes associated with response to cold, and seed composition-associated genes, such as SWEET transporter genes. Most of these SNPs had low MAF, suggesting that these variants have less profound deleterious effects in breeding programs.

### Population structure and diversity

The USDA germplasm collection accessions were classified into six subpopulations, most of which could be linked to geographic origins (**Supplementary Table S1**). Four major clusters were identified including modern varieties of diverse origins from different countries, global landraces collection, breeding lines and *G. soja* lines (**Figure 1D; Supplementary Figure S1**). We estimated the LD decay across three subpopulations viz landraces, cultivars and wilds to assess the frequency of recombination (**Supplementary Figure S2**). Wild population lines showed the fastest LD decay, dropping rapidly from ∼0.4 to below 0.1 within the first 50 Kb and reaching near-baseline levels (∼0.05) by 100-150 Kb, indicating higher recombination rates and genetic diversity. The landrace population exhibited intermediate LD decay starting around 0.5 and declining more gradually than the wilds. They maintained moderate LD (∼0.15-0.2) even at 300 Kb, showing a balanced genetic structure between wild and cultivated collections. The cultivar population, on the other hand, exhibited the slowest LD decay beginning at the highest (∼0.6) and maintaining elevated LD throughout, still showing substantial LD (∼0.2) at 300 Kb distance indicating extensive linkage blocks and reduced recombination.

### Haplotype map and blocks structure in the wilds, landraces and cultivars sub-populations

Based on the LD-based haplotype analysis, we observed haplotype blocks ranging from 4,378 to 20,874 among the wilds, cultivars and landraces populations (**Supplementary Figure S3**). The mean block size was 18.15, 31.10 and 45.36 kb, respectively, in the wilds, cultivars and landraces. Similarly, the mean SNPs per block were maximum in cultivars (45.4 SNPs/block) and least amongst the wilds (18.2 SNPs/block). Total coverage ranged from 79.5 to 696.4 Mb moving from wilds to cultivars. The haplotype block sharing revealed that the blocks most shared were between the cultivars and landraces (9,825), followed by landraces and *G. soja* (1,223) and the least between cultivars and *G. soja* (582).

### Tag SNPs analysis

Tagged SNPs represent a set of SNPs loosely linked to other SNPs within the LD block. The tagged SNPs analysis resulted in the identification of ∼1.3 million genome-wide SNPs (**Supplementary Figure S4**). The tag SNPs exhibited an even distribution across common (>20%), intermediate (10-20%) and low frequency (5-10%) variants, with roughly 350K-480K SNPs in each major category. Despite LD-based pruning typically removing rare variants, the analysis retained 519 rare variants (1-5% MAF), indicating a good representation of less frequent alleles that still contribute to haplotype diversity within the population (**Supplementary Figure S5**). Only 3 SNPs with MAF <1% were retained, which is expected since very rare variants are often population-specific and don’t serve as effective tags across diverse populations. The SNPs distribution suggests that the LD pruning successfully identified a comprehensive set of tag SNPs that capture both common population-wide variation and intermediate-frequency variants important for population structure and association studies. With ∼1.3 million total tag SNPs spanning the full MAF spectrum, this dataset should be well-powered for genome-wide association study (GWAS), population genetics and genomic prediction applications for soybean.

### GWAS analysis with Global HapMap-II data

Low-density SNP marker arrays have been typically deployed by soybean researchers to study the genetic control of variations in important traits. Since the release of the first soybean genome sequence, the density of publicly available markers for association panels has grown from 3K microsatellite markers to microarray-based markers to hundreds to thousands and tens of millions of SNPs generated via whole genome resequencing of large germplasm collections. Several QTLs have been mapped for important traits including the seed composition traits and have resulted in candidate genes for selection in breeding. The utilization of high-density SNP arrays presents an advantageous framework for the re-evaluation and refined mapping of QTLs through integration with previously acquired phenotypic datasets.

We employed the phenotypic data for seed protein content, oil, stem growth habit, lodging, and palmitic acid obtained from GRIN database for lines part of the GmHapMap-II collection (**Supplementary Table S7**). A genome-wide association study conducted with the new HapMap data identified two QTLs on Chr 20 and Chr 15 for seed protein content (**Figure 2A**). We deployed the SoySNP50K as the genotyping array, using the same phenotypic data, to investigate the effect of genotype marker density on detecting the causal genomic segment and underlying genes associated with these traits (**Figure 2B**). For seed protein content, we detected an additional associated genomic segment on Chr 15 with the HapMap array not detected by the SoySNP50K array (**Supplementary Table S8**). The lead SNP on 3,888,132 bp (MAF=0.18) contained two alleles “G” and “A” with the former allele leading to higher protein content and the latter allele for lower content (**Figure 2C**). We deployed the LD based approach to narrow down the genomic region to 3.22 Kb which contained a total of 20 SNPs (3,885,344 - 3,888,571 bp; r2>=0.85) and investigated this region for candidate genes mapping the gene with function sugar transporter SWEET (Glyma.15G049200; Wm8a.5 ID - Wm82_49162) within the linkage block identified on Chr 15 (**Figure 2D; Supplementary Table S8**). Similarly, for oil, we mapped the same genomic region of 3.22 Kb (3,885,344-3,888,571 bp; r2>=0.85) on Chr 15 for association with oil as mapped for protein content (**Supplementary Figure S6; Supplementary Table S9**). The lead SNP 3,888,099 bp (MAF=0.18) was part of the same LD block as crude protein content and harbored two alleles “T” and “C”. The gene expression analysis of several soybean samples revealed that this gene was specifically expressed in the seed coat and seed tissues, with relatively little expression in other major tissues (**Figure 2E**). Overall, the newly developed HapMap increases the power to detect causal genes at higher marker resolution.

**Figure 2:**
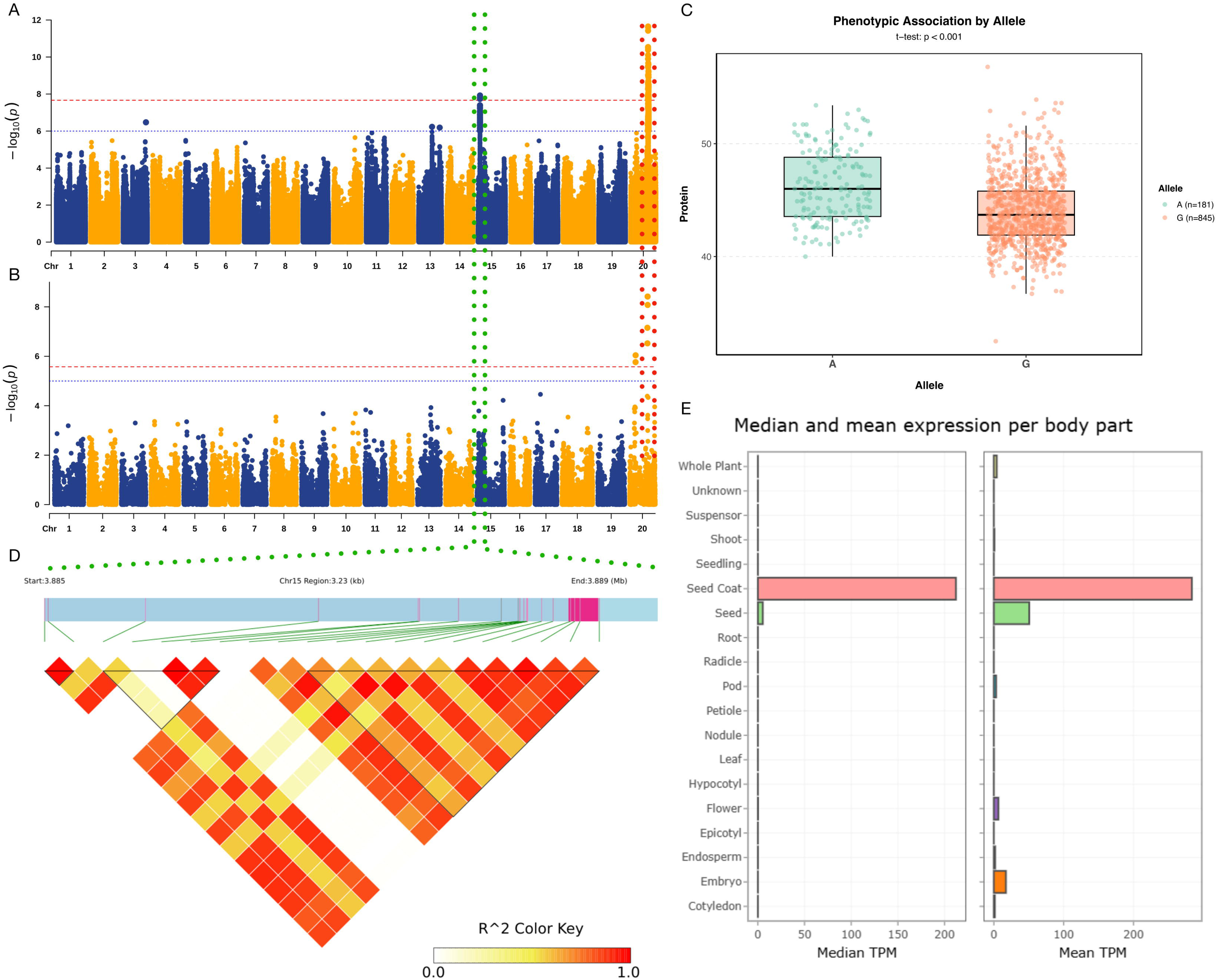
Association analysis of seed protein content in a soybean population. (A) Manhattan plot of GWAS analysis with haplotype map data, detecting genomic segments on Chromosomes 20 and 15. (B) Manhattan plot of GWAS analysis with the Soy50KSNP data detecting QTL on Chr 20. (C) Phenotypic distribution of two alleles for the peak signal detected on Chr 15 with haplotype map data. (D) Local LD heatmap of the QTL interval on chromosome 15, highlighting a candidate SWEET transporter gene. (E) Expression profiles of the candidate gene across diverse soybean samples.

GWAS analysis was also conducted for other important domestication-associated traits, such as lodging, stem growth habit and seed composition trait palmitic acid. A major QTL region on Chr 19 was identified for lodging resistance with the lead SNP anchored on 48,319,877 bp and two alleles A and C were responsible for lodging and no-lodging, respectively (**Supplementary Figure S7; Supplementary Table S10**). LD-based analysis revealed a region harboring nine candidate genes (Gm_Wm82_54928 - Gm_Wm82_54936) with functional annotation as flowering locus protein, F-box/LRR plant protein and iron-sulfur cluster assembly protein, etc. For the palmitic acid, we identified a major effect QTL on Chr 5 with the lead SNP position of 42,366,731 bp (**Supplementary Figure S8; Supplementary Table S11**). For stem growth habit, we identified a major QTL region on Chr 19 and LD- based candidate region revealed two genes harbored in the associated QTL region (**Supplementary Figure S9; Supplementary Table S12**).

### Superior haplotypes for protein content

We investigated the genic variations for gene *Glyma.15G049200* to analyse the haplotype patterns across the germplasm collection. A total of seven polymorphic sites were identified within the gene region, including one 3-bp InDel (3,888,412 bp) and multiple SNPs. Notably, one SNP (3,888,459 bp; G->A) was located in the coding region. Haplotype analysis revealed eleven distinct haplotypes, though five (H1-H5) accounted for majority of the variations (**Supplementary Table S13**). The dominant haplotype H1 was present in 822 lines, while H2 occurred in 173 lines (**Figure 3A**). The remaining haplotypes were considerably rarer, with H3 found in 53 lines, H4 in 12 lines, and H5 in 9 lines. Haplotypes H6-H11 were each present in fewer than 5 accessions.

**Figure 3:**
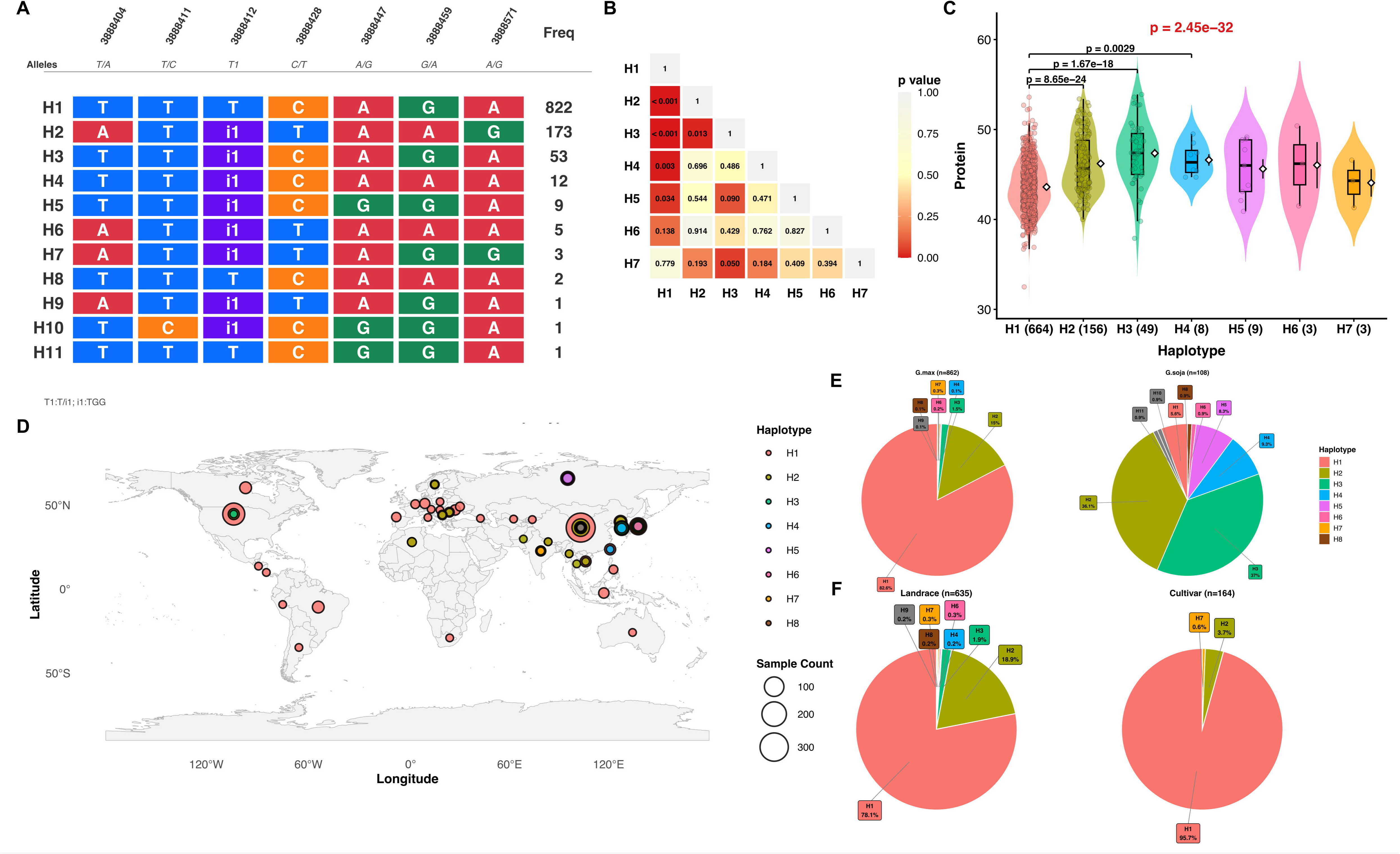
Superior haplotypes for seed protein content. (A) Composition of haplotypes and their distribution in the population. (B) Correlation of haplotypes (C). Association of haplotypes with the phenotypic spectrum. (D) Distribution of haplotypes globally in geographic locations. (E) Composition of haplotypes in two species: cultivated and *G. soja*. (F) Composition of haplotypes in different ecotypes: landraces and cultivars.

To identify functional variants, we tested for associations between haplotypes and protein content. This analysis revealed striking phenotypic differences among haplotypes. H2 showed significantly elevated protein levels compared to H1 (p < 2.22e-16), with H3 exhibiting intermediate values (**Figure 3B, 3C; Supplementary Table S14**). The protein content differences followed a clear gradient, with H1 associated with the lowest values, H2 and H3 with progressively higher levels, and the rare haplotypes showing variable patterns. We also examined oil content for the same haplotypes and found that H1 corresponded with higher oil levels while H2 showed reduced oil content, consistent with the well-documented negative correlation between seed protein and oil at this locus (Fleige et al., 2022; Goettel et al., 2023; **Supplementary Figure S10**). Geographic distribution of haplotypes revealed strong regional patterns. Chinese germplasm (n=418) displayed the greatest haplotype diversity, harboring H1 through H8, though H1 remained predominant at 76.8% (**Supplementary Figure S11**). In contrast, U.S. accessions (n=156) were nearly fixed for H1 (94.2%), with only a minor representation of H2 followed by H3. Japanese and South Korean materials showed intermediate diversity levels, with H1 frequencies of 55.9% and 41.7%, respectively, and substantial representation of H2 and H3. Several smaller collections from Brazil, Canada, Taiwan, Indonesia, Moldova, and Vietnam were completely fixed for H1 (**Figure 3D**). This geographic structure suggests that different breeding programs or environmental conditions have maintained varying levels of diversity at this locus.

Ecotype comparisons revealed distinct selective histories. Cultivars (n=163) exhibited extreme genetic uniformity with 96.3% carrying H1 and only 3.7% retaining H2 with a clear signature of breeding bottleneck. Landraces (n=632) maintained moderate diversity with H1 at 78.5% and H2 at 19%, plus low frequencies of H3 and the rare haplotypes H6 and H7. Wild soybean (*G. soja*, n=104) exhibited dramatically different haplotype structure, with nearly balanced frequencies of H1 (5.8%), H2 (37.5%), and H3 (38.5%), plus appreciable frequencies of H4 (9.6%) and H5 (8.7%) (**Figure 3E**). The absence of H3, H4, and H5 from cultivated materials and their presence in wild populations strongly suggests these represent ancestral variants that were lost during domestication. The species-level comparison reinforced these patterns. Cultivated soybean (*G. max*, n=859) was dominated by H1 (82.9%) with H2 at 15%, while wild soybean retained all five major haplotypes with a more even distribution (**Figure 3F**). This progression from balanced haplotype frequencies in wild populations, through moderate H1 dominance in landraces, to near fixation in modern cultivars illustrates the stepwise genetic erosion typical of crop domestication and modern breeding. The reduction in haplotype diversity represents approximately an 85% loss from wild to elite cultivated germplasm, with the initial domestication bottleneck eliminating H3-H5 and subsequent breeding selection driving H1 toward fixation.

### Additive Effects of Two QTLs Control Seed Protein and Oil Content

Two major QTLs on Chrs 15 and 20 were identified controlling seed protein and oil content with additive effects. Analysis of homozygous genotypes (n=80-671 per class) revealed that Chr 20 is the primary QTL, contributing +2.55% to protein content per Alt/Alt genotype, while Chr 15 acts as a minor modifier with a +0.65% effect (**Figure 4A** and **4B**). Both QTLs exhibited a strong negative correlation between protein and oil, with each 1% increase in protein associated with approximately 2% decrease in oil content. The reference genotype (Ref/Ref at both loci, 70% of germplasm) produced the highest oil content (18.3%) with the lowest protein (43.8%), while the double alternate genotype (Alt/Alt at both loci, 8% of germplasm) showed the inverse pattern with 47.0% protein and 12.3% oil. Two intermediate genotypes (Chr 15=Alt/Alt × Chr 20=Ref/Ref and Chr 15=Ref/Ref × Chr 20=Alt/Alt) offered balanced seed composition with 44.7-46.6% protein and 14.9-16.8% oil, representing viable targets for breeding programs requiring moderate improvements in both traits. A simple additive model (R²=0.135) accurately predicted protein content across all four homozygous genotype combinations, indicating these QTLs act independently and are suitable for marker-assisted selection (MAS) (**Figure 4C**). The substantial within-genotype variation suggests additional minor QTLs contribute to the observed phenotypic diversity in seed composition traits.

**Figure 4:**
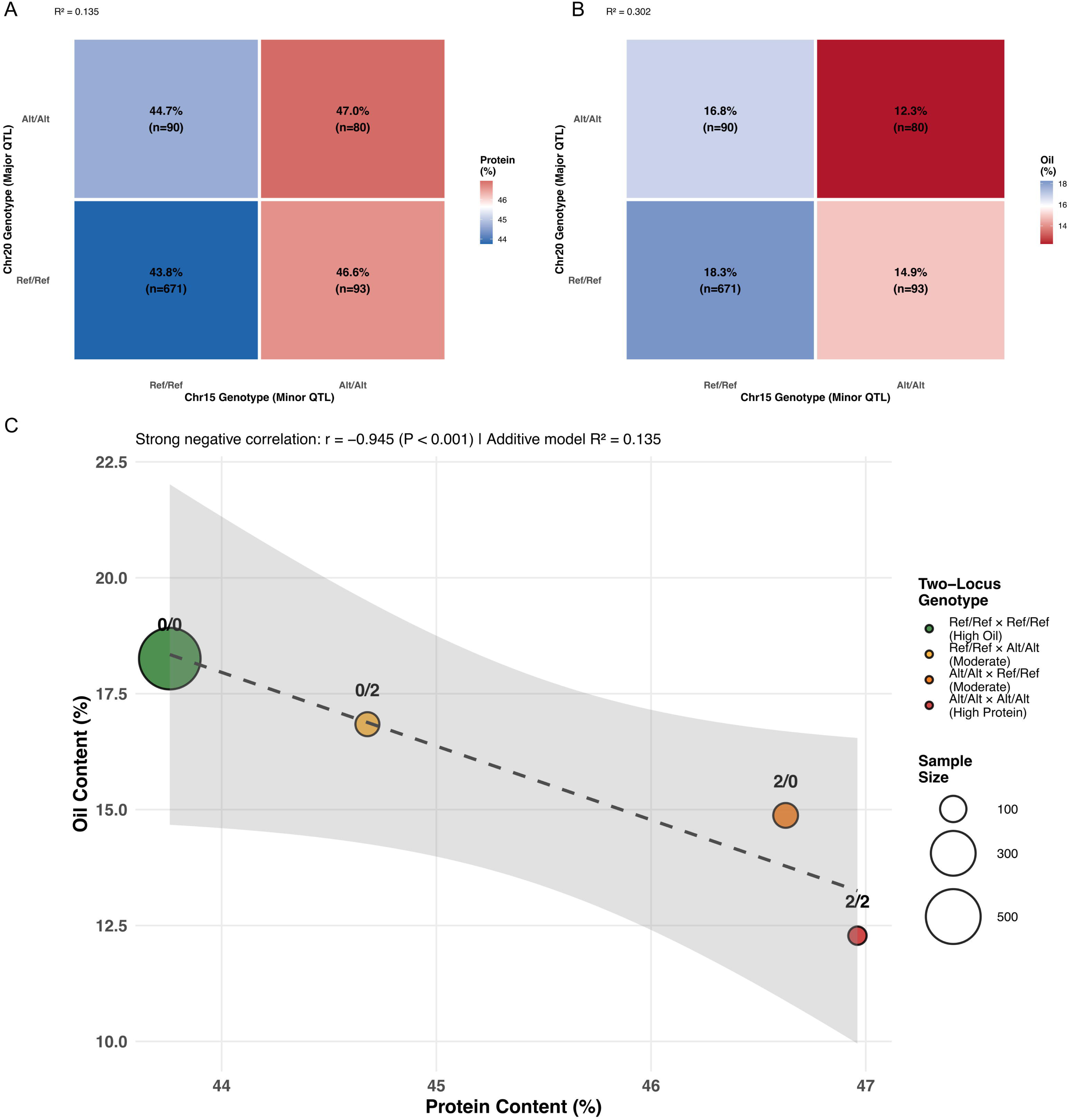
Two-locus additive model reveals independent QTL effects on soybean seed composition traits. **(A)** Protein content across four homozygous genotype classes showing additive effects of Chr15 and Chr20. Color intensity represents protein levels; darker red indicates higher protein content. Values show mean protein percentage with sample sizes in parentheses. The additive model explains 13.5% of phenotypic variance (R² = 0.135). **(B)** Oil content across the same genotype classes demonstrating inverse relationship with protein. Color intensity represents oil levels; darker blue indicates higher oil content. The additive model explains 30.2% of phenotypic variance (R² = 0.302). **(C)** Scatter plot illustrates the strong negative correlation (r = −0.945, P < 0.001) between protein and oil content across genotype classes. Bubble size indicates sample size (n = 80-671). Each point represents the mean of homozygous individuals, with the dashed line showing linear regression fit and the shaded area representing 95% confidence interval. The two-locus additive model accounts for both QTL effects without significant epistasis (P > 0.05). Chr15 contributes +0.65% protein per Alt/Alt genotype, while Chr20 contributes +2.55% protein per Alt/Alt genotype, demonstrating Chr20 as the major determinant of seed composition.

### Large-scale germplasm scanning reveals novel haplotype and copy number variations for major SCN loci

To expand beyond the limited sample size of initial haplotype screens, we analyzed the *rhg1* locus across a broader collection of HapMap soybean accessions from the germplasm collection. Our analysis revealed four haplotypes at this locus, with H1 being the most prevalent variant (**Figure 5A; Supplementary Table 15**). Among the 611 accessions with available phenotype data, H1 (n=562) displayed a strong association with susceptibility to SCN race 3 (**Supplementary Figure S12**). The two less frequent haplotypes showed distinct phenotypic patterns: H2 (n=31) was strongly associated with the resistant phenotype (p=4.3e-09), while H3 (n=18) exhibited association with moderate resistance to SCN race 3 (p=3.1e-07) (**Supplementary Figure S12**). A fourth rare haplotype, H4, was detected at very low frequency in the landrace germplasm (**Figure 5A**). Based on diagnostic SNP profiles, H1 corresponds to rhg1-c (Williams 82-like, susceptible reference type), H2 to rhg1-b (PI88788-type, high copy number with nine diagnostic non-synonymous SNPs), H3 to rhg1-a (Peking-type, low copy number with distinct non-synonymous variants), and H4 likely represents a sub-variant of rhg1- b. Global haplotype composition showed strong dominance of H1 (rhg1-c) across all regions, representing the reference susceptible type (**Figure 5B**). Among the total accessions analyzed, H1 was present at frequencies ranging from 80.8% to 100% depending on geographic origin and germplasm type. China, representing the largest collection (n=434), exhibited the following distribution: H1 (93.3%), H2 (3.9%), and H3 (2.8%), a pattern reflecting the center of soybean domestication with baseline genetic diversity in resistance alleles (**Supplementary Figure S13**). US germplasm (n=154) showed a notably different pattern with H1 at 86.4% and H2 at 13%, representing the highest frequency of H2 (rhg1-b) among all countries examined. The elevated H2 frequency represents a ∼3.3-fold enrichment compared to Chinese germplasm, likely reflecting decades of intensive breeding for SCN resistance. Japan (n=118) displayed haplotype diversity across all four variants, with H1 at 90.7%, H2 at 5.1%, H3 at 3.4%, and H4 at 0.8%. South Korea (n=110) exhibited the highest global frequency of H3 (rhg1-a) at 7.3%, accompanied by H1 at 86.4% and H2 at 6.4% (**Supplementary Figure S13**). This represents the most diverse haplotype composition among East Asian countries. In contrast, Russia (n=52) showed complete fixation for H1 (100%), with no alternative haplotypes detected.

**Figure 5:**
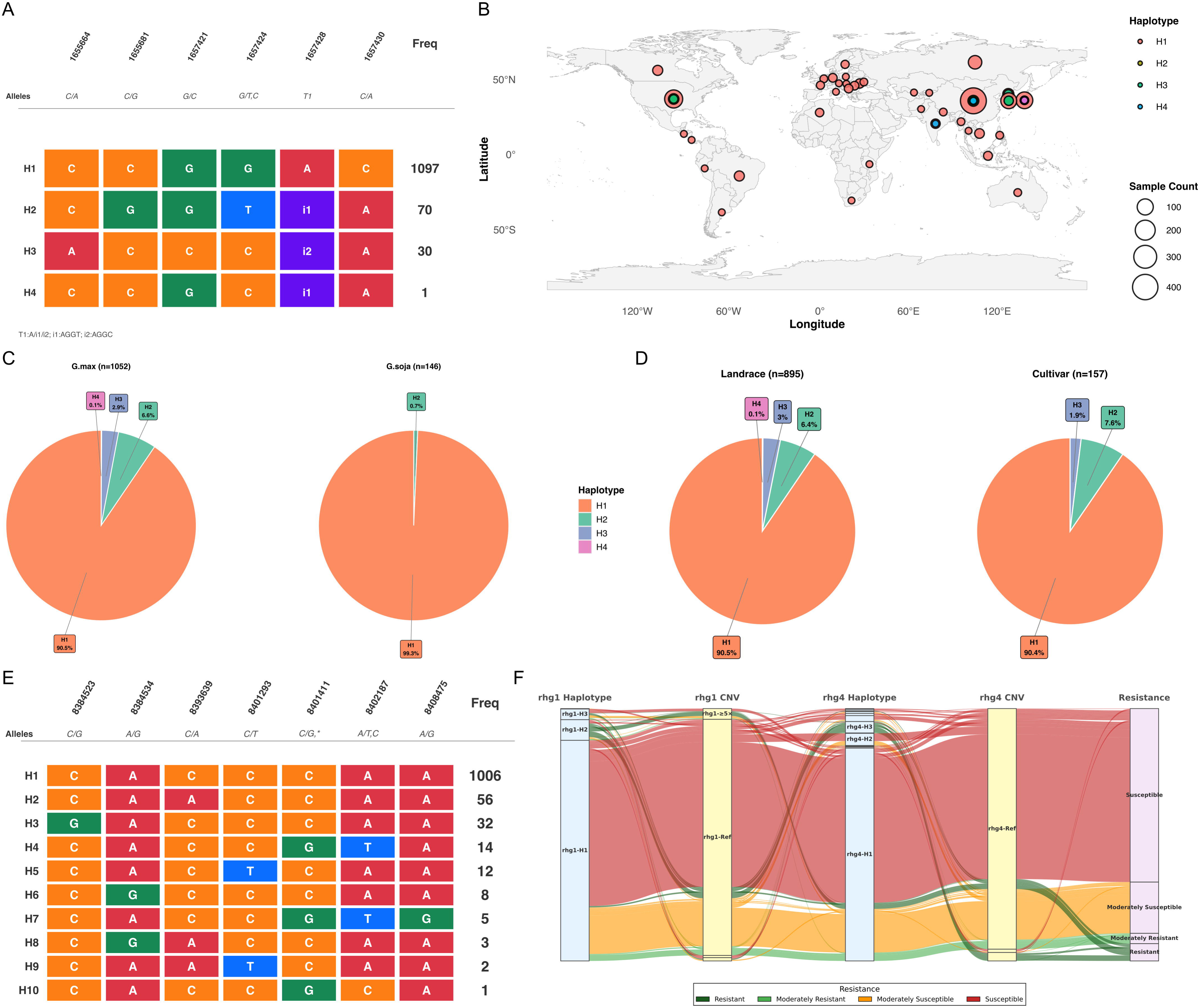
Haplotypes for SCN locus *rhg-1 and Rhg4* in germplasm collection. (A) Composition of haplotypes and their distribution in the population for *rhg-1* locus. (B). Distribution of *rhg-1* haplotypes globally in geographic locations. (C) Composition of haplotypes in two species: cultivated and *G. soja*. (D) Composition of haplotypes in different ecotypes: landraces, cultivars and wild-type. (E) Composition of haplotypes and their distribution in the population for Rhg4 locus. (F) Alluvial diagram illustrating the combinatorial effects of *rhg-1* and Rhg4 haplotypes and copy number variations on SCN race 3 resistance. The flow diagram visualizes the associations between genetic features at two major SCN resistance loci and phenotypic resistance outcomes. From left to right, the diagram shows: (1) rhg1 haplotype groups (rhg1-H1, rhg1-H2, rhg1-2-5×, and rhg1-Ref), (2) rhg1 copy number variation categories, (3) Rhg4 haplotype groups (rhg4-H1, rhg4-H2, and rhg4-Ref), (4) Rhg4 copy number variation categories, and (5) resistance phenotypes (Resistant, Moderately Resistant, Moderately Susceptible, and Susceptible).

Wild soybean (*G. soja*, n=146) exhibited near-complete fixation for H1 (rhg1-c) at 99.3%, with H2 detected at a very low frequency of 0.7% and complete absence of H3 and H4 haplotypes, indicating that resistance haplotypes are largely absent from the wild progenitor species. Cultivated soybean (*G. max*, n=1052) showed H1 at 90.5%, H2 at 6.6%, H3 at 2.9%, and H4 at 0.1%. The markedly lower frequency of alternative resistance haplotypes in wild germplasm compared to cultivated soybean suggests their predominantly post-domestication origin, either through spontaneous mutation during cultivation or selection of extremely rare variants not captured in the wild sample. Examination of haplotype frequencies across different germplasm categories revealed the impact of breeding selection on haplotype distribution (**Figure 5D**). Landrace accessions (n=895) displayed H1 at 90.5%, H2 at 6.4%, H3 at 3%, and H4 at 0.1%. This composition represents the baseline diversity present in traditional farmer-selected germplasm prior to the advent of modern systematic breeding. Modern cultivars (n=157) exhibited H1 at 90.4% and H2 at 7.6%, with H3 at 1.9% and a complete absence of H4. The H2 frequency in cultivars represents a ∼1.2-fold increase compared to landrace germplasm, indicating enrichment through modern breeding selection for SCN resistance. Across the cultivated germplasm collection, approximately 9-10% carried alternative resistance haplotypes (H2, H3, or H4), while ∼90% retained the susceptible H1 (rhg1-c) reference type.

The haplotype analysis of another major SCN locus rhg4 revealed the presence of seven SNPs affecting missense variations. The germplasm collection of a larger set of lines revealed additional variations than reported in our previous analysis (Patil et al., 2019). The haplotype analysis of the rhg4 gene showed the presence of 10 haplotypes, with two haplotypes dominant in the germplasm collection lines. H1 haplotype was observed in 1,006 lines followed by H2 in 56 lines and H3 in 32 lines, while the remaining 7 haplotypes (H4 - H10) were detected at very low frequencies (1-14 accessions each) and are collectively referred to as rare haplotypes. H4 was observed in 14 accessions, H5 in 12 accessions, and H6 in 8 accessions, while the remaining rare haplotypes (H7-H10) were each represented by 1-5 accessions. We examined the CNVs spectrum in the germplasm collection lines. Overall, we identified a total of 177 lines exhibiting CN variations with 168 lines harboring more than one copies in rhg1 locus (duplication) and 9 lines showing less than one copy (deletion) in this genomic segment (**Supplementary Table S16**). We confirmed the CNVs in known lines such as PI 88788 that possessed more than eight copies (∼8.6 CN). The highest copies were observed for PI 598124 (∼10.4 CN) followed by PI 639740 (∼10.2 CN) and S12-2894 (∼10.1) all showing around 10 copies of this locus. The CNV analysis revealed 64 lines with more than 5 copies of rhg1 locus and 95 lines showed more than 2 but less than 5 copies. Further, in the rhg4 locus 66 lines exhibited CN variations, with 49 lines harboring more than one copy. To investigate the interplay between haplotype identity and copy number variation in determining SCN resistance, we performed an integrated analysis of rhg1 haplotypes and CNV status across 615 accessions with complete phenotypic data. The combined analysis revealed distinct patterns of dosage-dependent resistance that varied by haplotype type (**Figure 5E; Supplementary Table S17**). For the H2 haplotype (rhg1-b; PI88788-type resistance), copy number variation showed a clear dose-response relationship with resistance phenotype. Among accessions with adequate sample sizes (n≥10), those carrying high copy numbers (3-5x duplication, n=17) exhibited the strongest resistance, with 76.5% showing resistant phenotypes and an additional 11.8% displaying moderate resistance (**Supplementary Table S17**). Accessions with moderate duplication levels (2-3x, n=10) showed intermediate resistance rates of 60%, with 20% moderately resistant and 20% moderately susceptible (**Supplementary Table S17**). This demonstrates a clear copy number threshold effect, where H2 requires at least 2-3x duplication to confer resistance, with high resistance achieved at 3-5x copy numbers of rhg1. Lower duplication levels (<2x, n=3) showed substantially reduced resistance rates (33.3%), though the limited sample size precludes definitive conclusions for this category. In striking contrast, the predominant H1 haplotype (rhg1-c; Williams 82-type susceptible) maintained high susceptibility regardless of copy number status. Among 552 accessions carrying H1 with reference single-copy configuration, only 3.3% showed resistant phenotypes, with 76.1% being completely susceptible. Even the small number of accessions with H1 duplications or deletions failed to confer resistance, indicating that the susceptible amino acid sequence cannot be compensated for by increased gene dosage. This demonstrates that specific amino acid polymorphisms in the GmSNAP18 protein, rather than expression level alone, are critical for resistance function.

The H3 haplotype (rhg1-a; Peking-type) showed distinct behavior from H2, with generally lower resistance rates even in the presence of CNV. Accessions with H3 and moderate duplication (2-3x, n=4) showed only 25% resistance rates, substantially lower than the 60% observed for H2 at comparable copy numbers. Higher copy number classes of H3 (3-5x and ≥5x) exhibited 0% resistance, despite amplification, although these categories contained 7 and 4 accessions, respectively. This pattern suggests that H3-mediated resistance has different genetic requirements than H2, consistent with previous reports of epistatic dependence on Rhg4 genotype for Peking-type resistance. Rare rhg1 haplotypes (H4, n=1) were detected at very low frequency, insufficient for meaningful resistance pattern analysis. The single accession carrying H4 exhibited a susceptible phenotype, suggesting it does not confer strong resistance independently; however, its potential contribution in specific genetic backgrounds or in combination with other resistance loci cannot be excluded. The alluvial plot reinforces the critical copy number threshold for H2-mediated resistance (**Figure 5E**). The transition from <40% resistance at low duplication to >75% resistance at high copy numbers demonstrates a functional threshold between 2-3x and 3-5x duplication levels. This quantitative assessment across a large germplasm collection provides breeding-relevant benchmarks, such as H2-type resistance, which requires a minimum of 2-3x duplication for partial resistance and 3-5x duplication for robust, reliable resistance to SCN HG type 0 (Race 3). Molecular characterization of the four identified rhg1 haplotypes revealed the specific polymorphisms underlying functional differences (**Figure 5E**). H1 carries the reference sequence with no resistance-associated substitutions. H2 is defined by characteristic SNPs at positions 1655664 (C → G), 1657421 (G → T), and 1657428 (T → C), corresponding to the PI 88788-type amino acid changes. H3 exhibits distinct polymorphisms, including substitutions at 1653312 (C → T) and 1655681 (C → A), as well as specific indel patterns (T2: A → i1, with an AGGT insertion), which define the Peking-type resistance. H4 carries a unique polymorphism combination not previously characterized, representing a rare allelic variant present at a frequency of 0.1%. The alluvial plot for SCN Race 5 revealed a strikingly contrasting pattern compared to Race 3 (**Supplementary Figure S14**). Regardless of rhg1 haplotype identity or copy number status, virtually all accessions were classified as susceptible or moderately susceptible, with no resistant phenotypes observed across any haplotype-CNV combination at either locus. This indicates that neither the PI88788-type (H2) nor the Peking-type (H3) rhg1 haplotypes, even in combination with rhg4 alternative haplotypes, confer effective resistance to SCN Race 5 in the germplasm collection examined, highlighting the race-specific nature of rhg1/rhg4-mediated resistance and the need to identify novel resistance sources for Race 5 management.

### Multi-class variations-based machine learning predicts SCN phenotypes in germplasm collection

The haplotype of gene sequences across the germplasm collection serves as a powerful tool for predicting phenotypes using machine learning approaches. We deployed a machine learning framework genomic prediction model that integrates multiple classes of variations including haplotype and CNVs since the SCN mechanism shows an association of CNVs in the *rhg1* locus in the soybean genome. Using an ensemble approach, we assessed several machine learning methods such as random forest, SVM, LightGBM and XGBoost to predict the phenotypes of lines with no phenotype based on the haplotypes of three major SCN-associated genes. We analysed the haplotypes of *rhg1* gene and used allelic composition to predict the SCN phenotypes for lines with no phenotypic data. For the *rhg1* gene we identified 9 high effect variants, such as missense variations and in frame disruptive insertion variant within the gene boundaries (Chr 18:1653419-1657430) and the gene-centric haplotype analysis resulted in identification of 4 major haplotype groups (**Figure 6A**). The major haplotypes Hap1 (65.15%) and Hap2 (12.88%) were found across 909 samples. We evaluated machine learning approaches for predicting SCN Race 3 resistance using rhg1 haplotype assignments and copy number variation data. For multi-class classification (R, MR, MS, S), our best model achieved a cross-validation accuracy of 73.1% using AdaBoost with haplotype features (**Figure 6B**). Machine learning-based genomic prediction was performed to assess the utility of haplotype and copy number variation (CNV) data in classifying soybean cyst nematode (SCN) race 3 resistance phenotypes (**Supplementary Table S18; Supplementary Table S19**). For multi-class classification (distinguishing resistance classes), combined haplotype and CNV features achieved the highest cross-validation accuracy of 0.724 ± 0.020 using AdaBoost, with an F1 score of 0.642. CNV features alone (0.727 ± 0.024, Gaussian NB) and haplotype features alone (0.723 ± 0.022, Gradient Boosting) showed comparable performance. Binary classification (resistant vs. susceptible) demonstrated substantially improved prediction accuracy, with combined features achieving 0.938 ± 0.032 using LDA and an F1 score of 0.937 (**Figure 6C**). CNV-based models performed exceptionally well in binary classification (0.932 ± 0.028, AdaBoost), while haplotype-based models also showed strong performance (0.929 ± 0.023, LDA). Across all feature sets, ensemble methods (Gradient Boosting, AdaBoost, Random Forest) and LDA consistently ranked among the top-performing algorithms, while simpler models like Logistic Regression showed moderate performance. Our approach demonstrates the utility of combining haplotype information with CN variations to predict the phenotype with higher accuracy.

**Figure 6:**
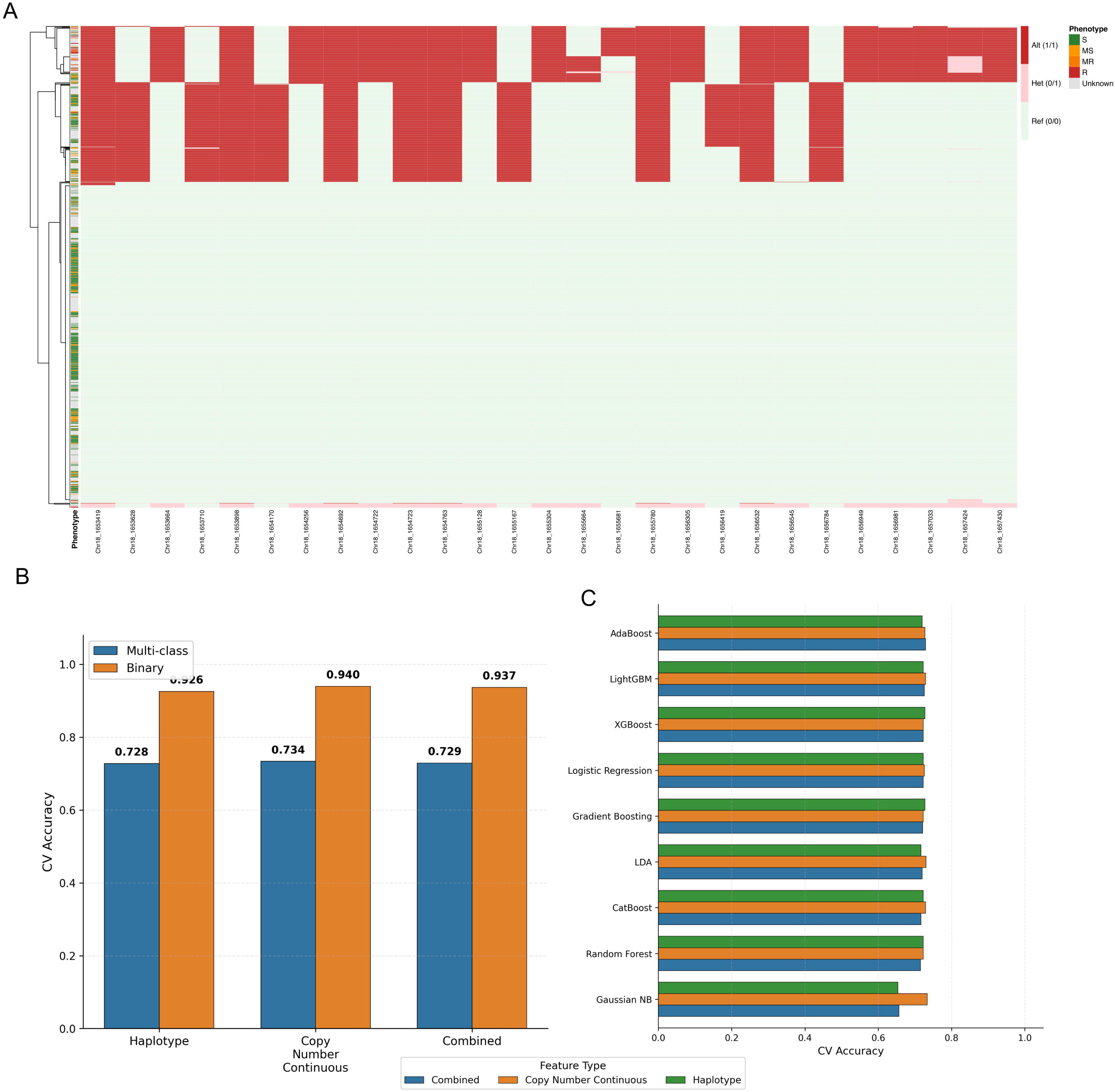
Machine learning-based genomic prediction of SCN resistance integrating haplotype and copy number variation data at the *rhg1* locus. **(A)** Heatmap showing phenotypic responses of soybean accessions to different SCN HG types (races 1, 3, 5, and 14). Accessions are hierarchically clustered (left dendrogram) with population structure indicated by colored bars (left margin). **(B)** Cross-validation accuracy comparison across feature types for multi-class (blue bars) and binary (orange bars) classification of SCN resistance. Three feature sets were evaluated: haplotype information alone, copy number variation data as continuous variables, and combined haplotype + CNV features. **(C)** Performance comparison of nine machine learning algorithms for multi-class SCN resistance prediction using combined haplotype and CNV features. Top-performing models include AdaBoost, LDA, and Random Forest.

### Deleterious alleles in soybean

Deleterious alleles have accumulated in the germplasm due to selection pressure. The analysis of germplasm collections can shed light on how the accumulation of deleterious alleles has occurred in different subpopulations of the germplasm collection. To evaluate how human selection and breeding shaped the genetic load in soybean (*G. max*), we quantified deleterious allele burden across three population groups representing key stages of soybean history: wild relatives (*G. soja*), traditional landraces, and modern cultivars. Deleterious mutations were identified using amino acid functional impact prediction with the Sorting Intolerant From Tolerant (SIFT) algorithm; nonsynonymous coding variants with SIFT scores < 0.05 were classified as deleterious, while those with SIFT ≥ 0.05 were classified as tolerated, paralleling approaches established in cassava and maize. Wild accessions carried the highest individual mutational load, averaging ∼12,500 deleterious alleles per individual, with substantial variance and numerous extreme outliers reaching nearly 15,000 alleles (**Figure 7A**). Landrace accessions showed a marked reduction, averaging approximately ∼7,250 deleterious alleles per individual, while modern cultivars exhibited the lowest burden, centered around ∼5,500 alleles per individual. Outlier cultivars with very low burdens (<2,500 alleles) likely reflect strong directional selection or genetic bottlenecks during elite line development. These three-way differences were all highly significant (Mann-Whitney U test; Wild vs. Landrace: p = 2.5×10^−76^; Wild vs. Cultivar: p = 7.1×10^−63^; Landrace vs. Cultivar: p = 1.6×10^−92^), confirming that both domestication and subsequent breeding improvement have progressively and significantly purged the soybean genome of deleterious alleles (**Figure 7B**).

**Figure 7.**
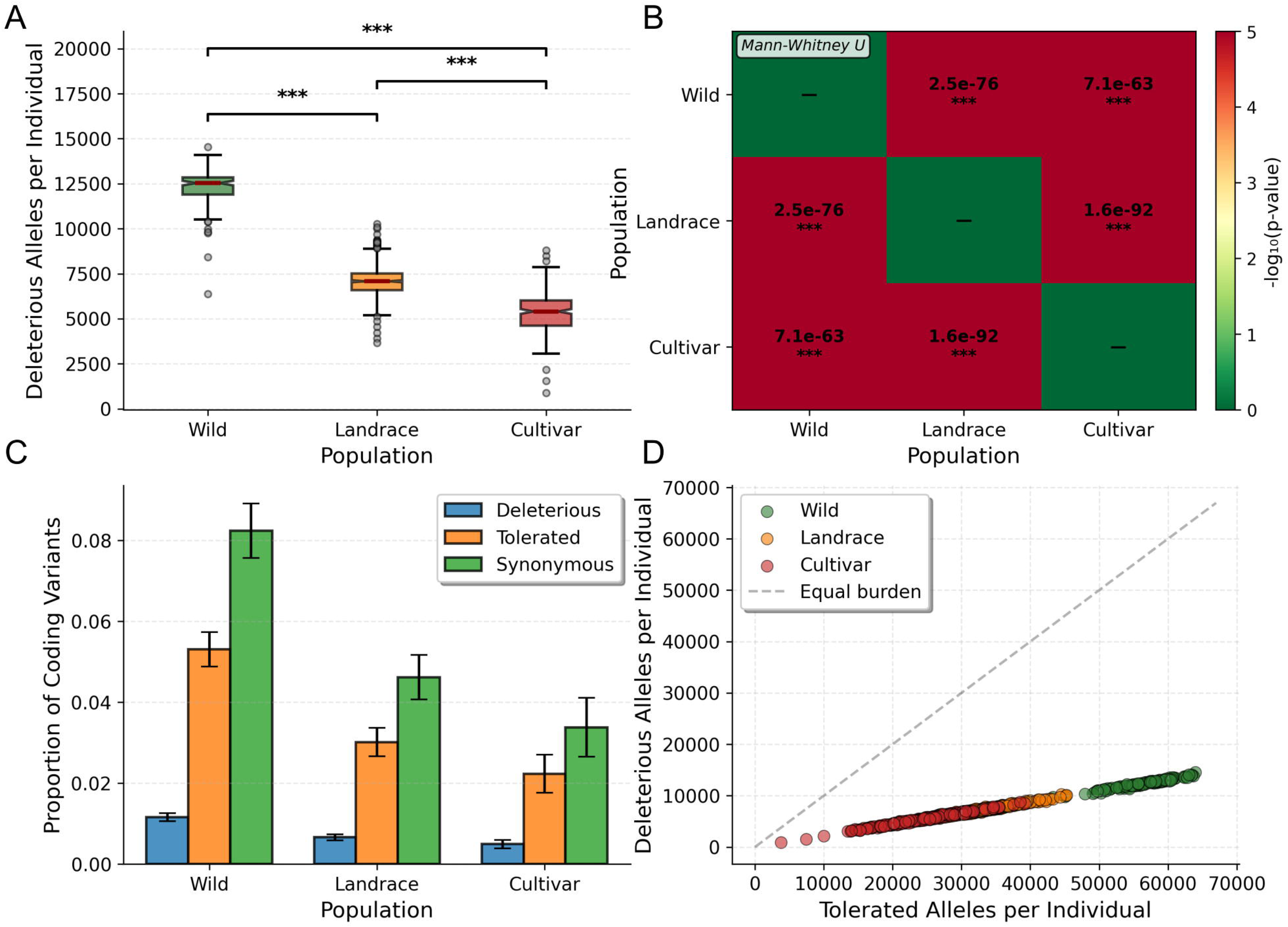
Deleterious mutation burden across soybean populations. (A) Box plots showing the number of deleterious alleles per individual in wild (*G. soja*), landrace, and modern cultivar accessions. Boxes represent the interquartile range (IQR); whiskers extend to 1.5× IQR; dots represent outliers. *** p < 0.001 (Mann-Whitney U test). (B) Pairwise Mann-Whitney U test p-value matrix (displayed as −log (p-value)) for all three population comparisons. Red indicates high significance. (C) Bar plots showing the proportion of coding variants classified as deleterious (nonsynonymous, SIFT < 0.05), tolerated (nonsynonymous, SIFT ≥ 0.05), or synonymous in each population. Error bars represent ± standard error. (D) Scatter plot of deleterious alleles vs. tolerated alleles per individual accession. The dashed line represents equal burden (slope = 1). Colors correspond to population groups as indicated in the legend.

To dissect the composition of genetic variation, we estimated the proportion of coding variants classified as deleterious (nonsynonymous, SIFT < 0.05), tolerated nonsynonymous (SIFT ≥ 0.05), and synonymous across all three population groups (**Figure 7C**). Across all categories, the proportion of each variant class consistently declined from wild to landrace to cultivar, reflecting the overall reduction in segregating diversity following domestication bottlenecks and breeding selection. Wild accessions harbored the highest proportion of all three categories: ∼1.3% deleterious, ∼5.3% tolerated, and ∼8.2% synonymous variants among coding sites. Landraces showed intermediate values (∼0.8% deleterious, ∼3.1% tolerated, ∼4.6% synonymous), and cultivars showed the lowest proportions (∼0.6% deleterious, ∼2.3% tolerated, ∼3.5% synonymous). Error bars indicate substantial within-group variation, consistent with population substructure and varying degrees of selection across accessions. We evaluated the relationship between deleterious and tolerated allele burden at the individual accession level (**Figure 7D**). Wild accessions (green circles) occupied the upper-right region of the plot, with tolerated allele counts ranging from ∼40,000 to ∼62,000 per individual and deleterious allele counts of ∼10,000-14,500. Landrace accessions (orange circles) formed an intermediate cluster, while cultivar accessions (red circles) were concentrated at the lower-left, with tolerated allele counts of ∼5,000-20,000 and deleterious counts of ∼1,000-8,000. Furthermore, understanding and utilizing the genetic resources of these wild accessions is of great significance for soybean genetic improvement and enhancing their environmental adaptability.

### Gene-centric haplotyping and a soybean haplotype database

Candidate gene discovery is an important component for enhancing important traits and towards developing improved varieties suited for ever-changing environmental conditions. SNPs that remain in high LD form a haplotype and do not show breakage due to recombination events staying together over generations. Analyzing these haplotypes and their association with traits of interest is a critical aspect of haplotype-based crop improvements. We performed the gene-centric haplotyping of all genes using “geneHapR” with more than two variants within the gene boundaries. Using this approach, we narrowed down to 46,441 such genes and performed gene-centric haplotype analysis. Several of these genes associated with protein, oil and seed composition traits harbored SNPs useful for identifying the beneficial alleles for trait improvement. A few examples of these genes include the sugar transporter gene, which contained seven major haplotypes with H1 distributed mainly among the landraces and H2 mainly observed in the *G. soja* lines.

Modern haplotype databases contain extensive genomic variation data, typically comprising millions of SNPs and thousands of small InDels across diverse soybean germplasm collections. These databases organize variations into distinct haplotype blocks, identifying 10,000-50,000+ haplotypes depending on the variations amongst the lines under study. Soybean haplotype database (SoyHapDB) harbors the haplotypes information that the users can extract interactively based on their search criteria (http://75.101.211.130/SoyHapDB/). Users can extract results through multiple approaches including gene-based searches, chromosomal region queries, and trait-associated variant mining. The database offers visualization tools for haplotype structure, LD patterns, and population genetics analyses. Interactive interfaces allow researchers to filter variants by allele frequency, functional impact, or specific germplasm subsets. These haplotype databases streamline haplotype analysis by providing pre-computed haplotype blocks, automated tag SNP selection, and integration with phenotypic data. Users can efficiently identify favorable haplotypes for breeding programs, perform GWAS, and trace allelic diversity across different soybean populations, significantly accelerating genomics-assisted breeding efforts.

## Discussion

Using whole-genome sequencing data from a large collection of 1,278 soybean accessions, we developed a global soybean haplotype map that provides a comprehensive resource for genetic studies and genomics-enabled breeding. Since the release of the first soybean reference genome, several high-quality soybean genomes have been furnished owing to long-read sequencing, demonstrating significant improvements in genome quality (Schmutz et al., 2010; Garg et al., 2023). Although several earlier variation and haplotype maps have been generated for soybean, these efforts primarily examined local germplasm panels and relied on older, less contiguous genome assemblies (Kim et al., 2021, Torkamaneh et al., 2021, Valliyodan et al., 2021, Yang et al., 2022, Liu et al., 2022). By assembling a diverse panel of 1,278 globally distributed cultivated and *G. soja* accessions and aligning all data to a modern high-quality genome, we produced a unified and high quality catalog of 11.37 million SNPs and 2.05 million InDels. Population structure analyses clearly separated domesticated and wild accessions and further resolved wild soybean into geographically defined clusters, consistent with patterns seen in other major crops. Linkage disequilibrium patterns confirmed reduced recombination in cultivated relative to wild soybean (Song et al., 2015), with wild accessions displaying rapid LD decay and contributing diversity absent from cultivated germplasm (Khan et al., 2024).

While low-density SNP arrays have facilitated trait mapping in soybean, these platforms lack coverage of insertion-deletion polymorphisms and structural variants, capturing only a fraction of the genomic variation landscape (Song et al., 2013). Whole-genome resequencing-derived variant matrices capture InDels and SVs undetectable by arrays, enabling complete characterization of allelic diversity at functionally important loci. Deploying the GmHapMap-II matrix led to the identification of a significant genomic region associated with crude protein content overlooked by low-density SNP array data, in addition to identifying the known major genomic signal on chromosome 20 (Diers et al., 1992; Marsh et al., 2023). The identification of extensive genetic variations between cultivated and wild species accessions provide insight into the domestication process and how breeders selected alleles amidst the presence of deleterious mutations within the germplasm. The Chr 15- and Chr 20- QTLs collectively explained 13.5% of protein variance, representing a substantial contribution for two loci in this polygenic trait. This is consistent with major QTLs in previous soybean seed composition studies, where individual loci typically explain 3-15% of phenotypic variance (Bandillo et al., 2015; Hwang et al., 2014). The additive nature of these QTLs facilitates marker-assisted breeding through independent pyramiding of favorable alleles. The superior haplotype analysis of the protein gene on Chr 15 shows strong evidence of directional selection favoring H1 with H1 carrying favorable alleles for protein content. The ∼20% retention of H2 in landraces suggests adaptive value in specific environments, while H3, unique to wild germplasm, underscores the importance of conserving wild soybean diversity. Our comprehensive analysis of 1,278 accessions corroborates the fundamental haplotype structure at Glyma.15G049200 (GmSWEET39) reported by Zhang et al. (2020). Our expanded panel of 11 major haplotypes, with H1 dominant in 822 (76%) cultivated accessions versus 335 (70.7%) in their study, validates the importance of this haplotype for marker-assisted selection.

Rhg-1 locus is a major genomic region conferring resistance to cyst nematodes with a combination of haplotype and CNV patterns (Cook et al., 2012). The three genes within each repeat GmSNAP18 (alpha-SNAP), an amino acid transporter, and other genes contribute to resistance in a dosage-dependent manner (Cook et al., 2014). Similarly, Rhg4 encodes a serine hydroxy methyltransferase (SHMT) and also exhibits CNV, with copy numbers ranging from 1 to 4.3 copies in resistant lines (Patil et al., 2019). The results of rhg-1 locus haplotype analysis demonstrate that SCN resistance haplotypes are rare globally, show strong geographic structuring, arose post-domestication and have undergone region-specific selection in modern breeding programs (Lee et al., 2015). The contrasting distribution patterns of H2 and H3 across geographic regions and germplasm types provide insights into breeding history and regional adaptation strategies for SCN management. Recently, our group demonstrated the identification of superior haplotypes/alleles for broad-spectrum SCN resistance from +1100 USDA germplasm using systematic data mining efforts such as haplotype, InDel and CNV analysis (Chhapekar et al. 2025). In this study, the comprehensive analysis of 1,278 accessions confirms the presence of multiple functional haplotypes at the rhg1 locus previously reported by Patil et al. (2019) in a diverse panel of 106 soybean lines. While previous study identified three major haplotype groups (rhg1-a, rhg1-b, and rhg1-c) based on nine amino acid substitutions, our expanded germplasm screen identified four distinct haplotypes (H1-H4) with similar functional associations. The consistency between studies with the susceptible haplotype dominating at approximately 91% and resistance haplotypes comprising 9-10% of germplasm validates the robustness of these haplotype classifications. However, our larger sample size enabled the discovery of a rare fourth haplotype (H4, 0.1% frequency) present exclusively in landrace germplasm, demonstrating that comprehensive germplasm screening continues to reveal cryptic allelic diversity that may be valuable for breeding. Our analysis of 1,191 globally distributed accessions revealed distinct geographic patterns of rhg1 haplotype distribution. The ∼3.3-fold enrichment of H2 (rhg1-b; PI88788-type) in U.S. germplasm (13%) compared to Chinese collections (3.9%) provides quantitative evidence of intensive selection pressure in major soybean-producing regions. This contrasts with the highest frequency of H3 (rhg1-a; Peking-type) observed in South Korea (7.3%), suggesting regional breeding priorities or differential nematode population pressures have shaped haplotype frequencies. Importantly, this study substantially expands on the CNV analysis of Patil et al. (2019), which characterized copy number variation in a panel of 106 lines. Our analysis across 1,191 accessions identified 184 lines with CNV at the rhg1 locus, including lines with exceptionally high copy numbers such as PI 598124 (∼10.4×), PI 639740 (∼10.2×), and S12-2894 (∼10.1×), well exceeding the range previously reported (Lee et al., 2015). Notably, copy number at Rhg1 has been shown to be dynamic even within released inbred cultivars, with individual plants of the same variety carrying between 9 and 11 copies, and selection for higher copy number directly improving SCN resistance (Lee et al., 2016). At the rhg4 locus, our expanded analysis revealed 14 haplotypes compared to the three reported previously, including several novel rare haplotype combinations not previously characterized. These unique haplotype-CNV combinations at both loci represent a substantially richer allelic landscape than previously documented and offer new targets for breeding programs seeking durable SCN resistance. This integrated haplotype-CNV analysis demonstrates that SCN resistance at the rhg1 locus results from the combined effects of specific amino acid-altering polymorphisms and gene copy number, with distinct functional requirements for different haplotype classes. The PI88788-type H2 can confer resistance through high copy number alone, while Peking-type H3 appears to require additional genetic factors, consistent with its known epistatic interaction with the Rhg4 locus (Cook et al., 2014). These findings provide quantitative guidelines for MAS, indicating that breeding for durable SCN resistance requires both selection for appropriate haplotype identity and verification of adequate copy number levels.

Genomic prediction has become essential in soybean breeding, with studies demonstrating prediction accuracies ranging from 0.3 - 0.7 for yield, seed composition, and agronomic traits depending on trait heritability and population structure (Duhnen et al., 2017; Zhang et al., 2016; Matei et al., 2018). Traditional genomic prediction models such as GBLUP and ridge regression BLUP assume additive effects of biallelic markers and have proven effective for many traits. However, these SNP-based approaches inadequately capture structural variations such as copy number variations (CNVs) that play critical roles in certain disease resistance traits. The rhg1 locus exemplifies this limitation. At rhg1, resistance to SCN is conferred by tandem repeats of a 31.2-kb segment containing multiple gene copies, with copy numbers ranging from 1 in susceptible genotypes to 10 in resistant lines (Cook et al., 2012; Cook et al., 2014). These structural variations cannot be adequately represented by binary SNP markers, as gene copy dosage directly influences resistance levels, with at least 5.6 copies of PI88788-type rhg1 required for effective resistance (Patil et al., 2019). Furthermore, rhg1 and Rhg4 exhibit race-specific epistatic interactions, with Peking-type resistance requiring both loci while PI88788-type resistance functions primarily through rhg1 alone (Yu et al., 2016). Machine learning approaches, including Random Forest, XGBoost, and neural networks, offer distinct advantages by modeling non-linear relationships and accommodating CNV dosage as continuous predictors (Montesinos-López et al., 2022; Gill et al., 2022). Our machine learning approach integrating both haplotype and CNV information at rhg1 represents a novel strategy for predicting SCN resistance. The substantial performance gap between binary and multi-class classification (>20% accuracy improvement) suggests that while genomic features effectively discriminate against broad resistance categories, fine-grained phenotypic classification remains challenging, potentially reflecting the quantitative and polygenic nature of SCN race 3 resistance or limitations in phenotypic resolution. The comparable performance of haplotype-only, CNV-only, and combined feature sets indicates that both genomic variation types capture largely overlapping resistance signals at known loci (rhg1/rhg4), though their combination provides marginal improvements in certain models. The superior performance of ensemble methods and LDA suggest non-linear relationships between genomic features and resistance phenotypes, with feature interactions playing important roles in prediction accuracy. The high binary classification accuracy (>93%) demonstrates the practical utility of this approach for resistance screening applications, potentially enabling marker-assisted selection strategies for SCN race 3 resistance breeding programs. Future work should validate predictions in independent germplasm panels, explore model performance across different SCN races, and investigate feature importance of identifying the most informative haplotype-CNV combinations.

We analyzed the mutation in the germplasm collection with haplotype map data and the interesting feature is that mutation burden was reduced in domesticated relative to wild soybean accessions. Unlike most domesticated crops where deleterious alleles accumulate following domestication (Sun et al., 2023; Valluru et al., 2019; Ramu et al., 2017), the progressive reduction in deleterious allele burden from wild *G. soja* through landraces to modern cultivars demonstrates that purifying selection has operated efficiently throughout soybean’s domestication and improvement trajectory. Unlike clonally propagated crops such as cassava, where limited recombination paradoxically increases deleterious burden following domestication, soybean’s obligate sexual reproduction has enabled recombination to continuously expose and purge recessive deleterious alleles. The disproportionate reduction in deleterious relative to synonymous variant proportions confirms that this burden reduction reflects genuine purifying selection rather than a passive consequence of reduced overall diversity. The GmHapMap-II is a significant resource for assessing the gene-centric haplotyping of important loci/genes linked with traits. Our haplotype database facilitates easy extraction of haplotype and SNP information, making this resource accessible to researchers with varying computational expertise.

This study presents a high-resolution haplotype map that significantly advances our understanding of genetic variation in the US soybean germplasm collection and its relationship to key agronomic traits. By integrating variant calling, quantitative genetics, structural variation analysis and machine learning approaches, we demonstrated the power of haplotype-based analyses for dissecting complex trait architecture in soybean. The identification of previously undetected minor-effect QTL and the detailed characterization of epistatic interactions at SCN resistance loci highlight the limitations of low-density marker platforms and underscore the value of whole-genome sequencing for germplasm characterization. Our machine learning framework for integrating haplotypes and CNVs at rhg1 provides a template for genomic prediction in regions with complex structural variation, which are often challenging to analyze with conventional approaches. The gene-centric haplotype database we developed serves as a valuable community resource, facilitating marker-assisted selection, genomic selection, and candidate gene identification for soybean improvement. Overall, this work establishes a foundation for haplotype-based breeding strategies, with insights into genetic mechanisms underlying quantitative trait variation applicable to both fundamental research and soybean improvement programs.

## Experimental procedures

### Sequencing data

This study utilized two soybean germplasm collections: (i) 1,116 accessions for which whole-genome sequencing data were publicly available, sourced from prior studies covering globally representative diverse accessions (Zhou et al., 2015), accessions from the USA (Valliyodan et al., 2016), Chinese germplasm (Fang et al., 2017), and parents of the soybean nested association mapping (NAM) population (Song et al., 2017); (ii) an additional 162 accessions that were newly sequenced as part of the present study. For DNA extraction, leaf tissue was collected and stored frozen prior to homogenization using a Qiagen TissueLyser. Approximately 100 mg of homogenized tissue was used for DNA isolation with the Qiagen Plant DNeasy Mini Kit according to the manufacturer’s recommended protocol. DNA concentration and purity were assessed using a NanoDrop spectrophotometer. For 142 of the newly sequenced accessions, paired-end sequencing libraries were prepared using the KAPA Hyper Prep Kit (Kapa Biosystems, Wilmington, MA, USA) in accordance with the manufacturer’s guidelines (KR0961 - v5.16). Short-read whole-genome resequencing was carried out on an Illumina HiSeq 2000 platform.

### Identification of variants

Sequencing reads from all 1,278 accessions were processed through a standardized bioinformatics workflow built around GATK4 to generate a comprehensive catalog of genomic variants. Raw paired-end reads of 100-150 bp length were first subjected to quality trimming using Trimmomatic v0.39, which removed low-quality reads and clipped bases falling below quality thresholds (Bolger et al., 2014). Read quality was evaluated using the Raspberry tool v0.2 (Katta et al., 2015). Quality-filtered reads were subsequently aligned to the *G. max* reference genome version 5 using BWA-MEM v0.7.19 (Li et al., 2013; Garg et al., 2023). Resulting SAM alignment files were converted, coordinate-sorted, and stored in BAM format using the Picard toolkit v3.0.0 (Picard toolkit, 2019). Per-sample variant detection was carried out using the HaplotypeCaller module of GATK v4.2.6.1 (McKenna et al., 2010). The resulting variant dataset was subsequently phased and missing genotypes were imputed using BEAGLE v5.1 (Browning and Browning, 2016). Functional annotation of identified variants was performed with SnpEff v3.0 (Cingolani et al., 2012). For phylogenetic reconstruction, genome-wide SNPs were pre-filtered using SNPhylo (Lee et al., 2014) with the parameters -m 0.01, -M 0.01, and -l 0.5 to reduce redundancy, and the retained SNPs were then used to infer a maximum-likelihood phylogeny with FastTree2 v2.1.10 SSE3 (Price et al., 2010) using the -nt -gtr settings.

### Phenotype collection and metadata

Phenotypic data for protein content, oil content, stem growth habit, lodging score, palmitic acid concentration, and SCN HG type reactions used in this study were sourced from the USDA Soybean Germplasm Collection general evaluation trials, which encompass morphological, agronomic, and seed composition datasets. These field-based evaluations were carried out across multiple locations stratified by maturity group, with some accessions evaluated over multiple growing seasons. Total protein and oil concentrations were quantified using methods appropriate to seed coat pigmentation. For accessions with yellow seed coats, protein and oil content were determined by near-infrared reflectance spectroscopy applied to whole seed samples. Accessions with dark or pigmented seed coats were analyzed separately, with total protein quantified via the Kjeldahl method and seed oil content measured using the Butt method (Bandillo et al., 2015).

### Genome-wide association analysis

Genome-wide association analysis was conducted using two complementary statistical approaches, the mixed linear model (MLM) and FarmCPU, both executed within the rMVP framework (Yin et al., 2021). Associations were tested for protein content, oil content, stem growth habit, lodging, and palmitic acid concentration using imputed genotype data across all accessions with available phenotypic records. To account for confounding due to population stratification, the first three principal components derived from genome-wide marker data were incorporated as covariates. The genome-wide significance threshold was set at a Bonferroni-corrected value of 0.05 divided by the total number of markers tested.

### Linkage disequilibrium and haplotype analyses

Linkage disequilibrium heatmaps were constructed using LDBlockShow (Dong et al., 2020), computing both R² and D’ statistics, with haplotype block boundaries defined using the Gabriel et al. (2002) confidence interval method for D’-based visualization. Input for the accompanying Manhattan plot was derived from FarmCPU-based protein GWAS results obtained from filtered, unimputed SNP data. Haplotype block structure was characterized using the confidence interval approach of Gabriel et al. (2002) as implemented in PLINK v1.9 (Purcell et al., 2007) with default settings. Pairwise R², D’, and allele frequency estimates were extracted for relevant genomic sites using PLINK v1.9, whereby sites in linkage disequilibrium with the GWAS-identified SNP were first identified using the --show-tags flag with a --tag-r2 threshold of 0.9, followed by generation of detailed pairwise statistics for the tagged sites using the --r2 dprime option. The gene-based haplotype analysis was performed with geneHapR package (Zhang et al., 2023). Functional impacts of nucleotide variants on proteins were assessed using SIFT 4G v.2.0.0, with variants scoring below 0.05 classified as putatively deleterious (Kumar et al., 2009).

### Multi-omics genomic prediction for SCN trait

Genomic prediction was performed using a multi-omics machine learning approach. Haplotype features were combined with structural variation (CNV) data to create a comprehensive feature matrix. Population stratification was corrected using principal component analysis. Model selection included seven machine learning algorithms with cross-validation to identify optimal predictors.

### Haplotype database development

The haplotype database development was performed using the Linux platform. We used Apache2 and Postgres for developing the tables and querying the database. The database queries were linked with the SNP ID across different tables stored in the database.

## Acknowledgment

We greatly acknowledge the funding support from the United Soybean Board (Grants #2312-209-0201 and #1320-532-5615) and the United States Department of Agriculture - National Institute of Food and Agriculture (USDA-NIFA) Award No. 2024-67013-42812). The computation for this work was performed on the high-performance computing infrastructure operated by Research Support Solutions in the Division of IT at the University of Missouri, Columbia MO https://doi.org/10.32469/10355/97710

## Author contributions

HTN oversaw the project and conceived the research idea; HTN and AWK designed the experiments and managed the project. AWK, DD, QS and SSC analyzed the genomic data. TDV and HY contributed to plant growth, sample preparation, DNA quality tests and DNA sequencing. AWK, TDV, SSC, VG, RKV and HTN wrote the manuscript. All authors discussed the results and contributed to the final manuscript.

## Data availability statement

All data supporting the findings of this study are available within the paper. The raw sequencing reads have been deposited to NCBI SRA with BioProject ID PRJNA1442058.

