## Supplementary Materials for "Next-Generation Soybean Haplotype Map as A Genomic Resource for Enhanced Trait Discovery and Functional Analysis"

**Supplementary Figures**

**Supplementary Figure S1: Phylogenetic tree for the population studied**

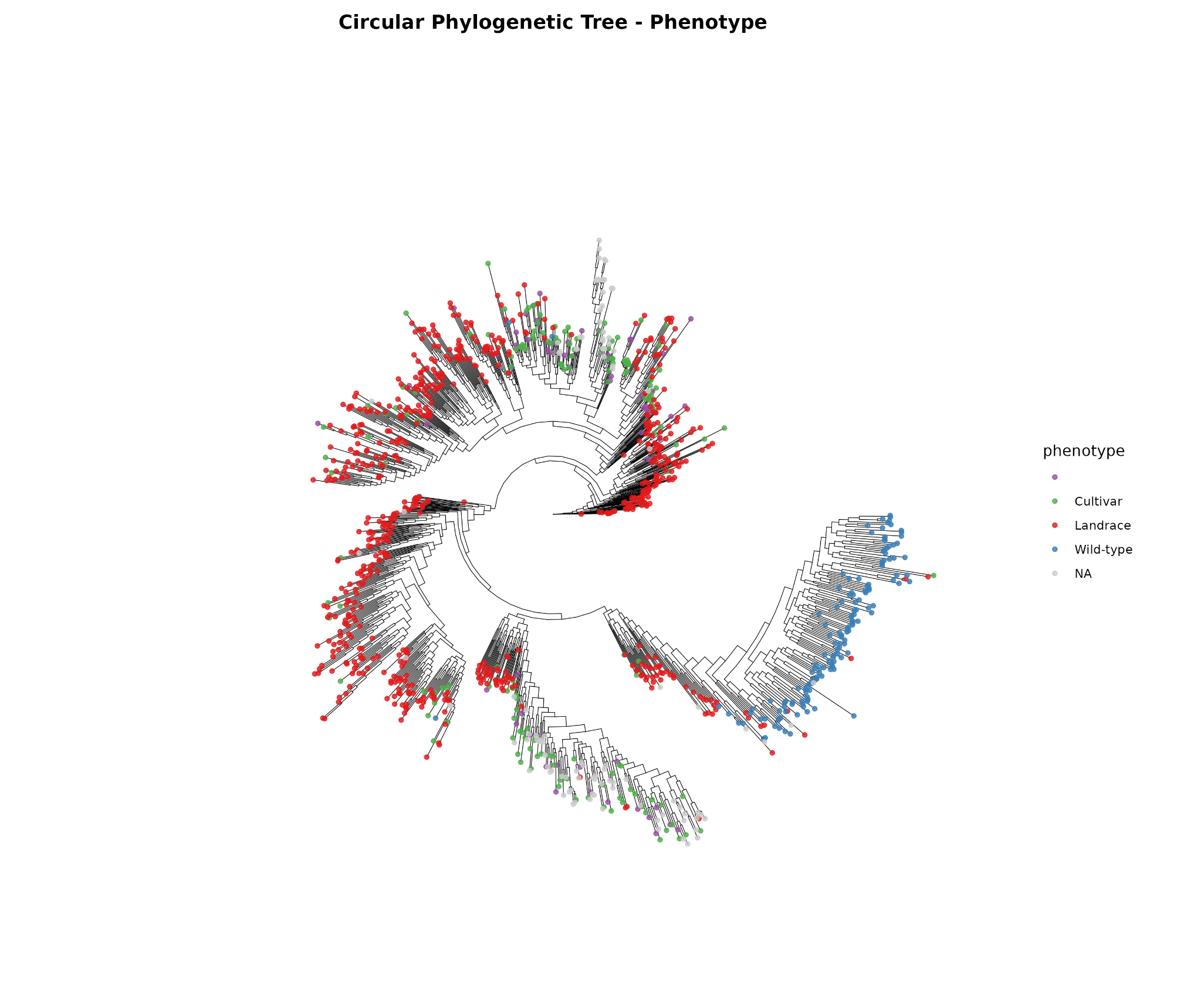

**Supplementary Figure S2: LD decay patterns across soybean subpopulations**

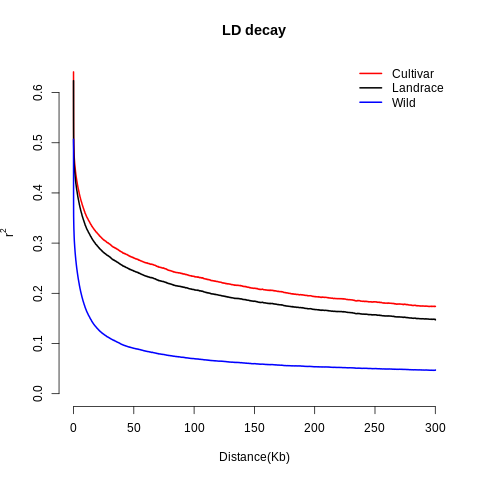

**Supplementary Figure S3: Haplotype blocks observed in different soybean subpopulations**

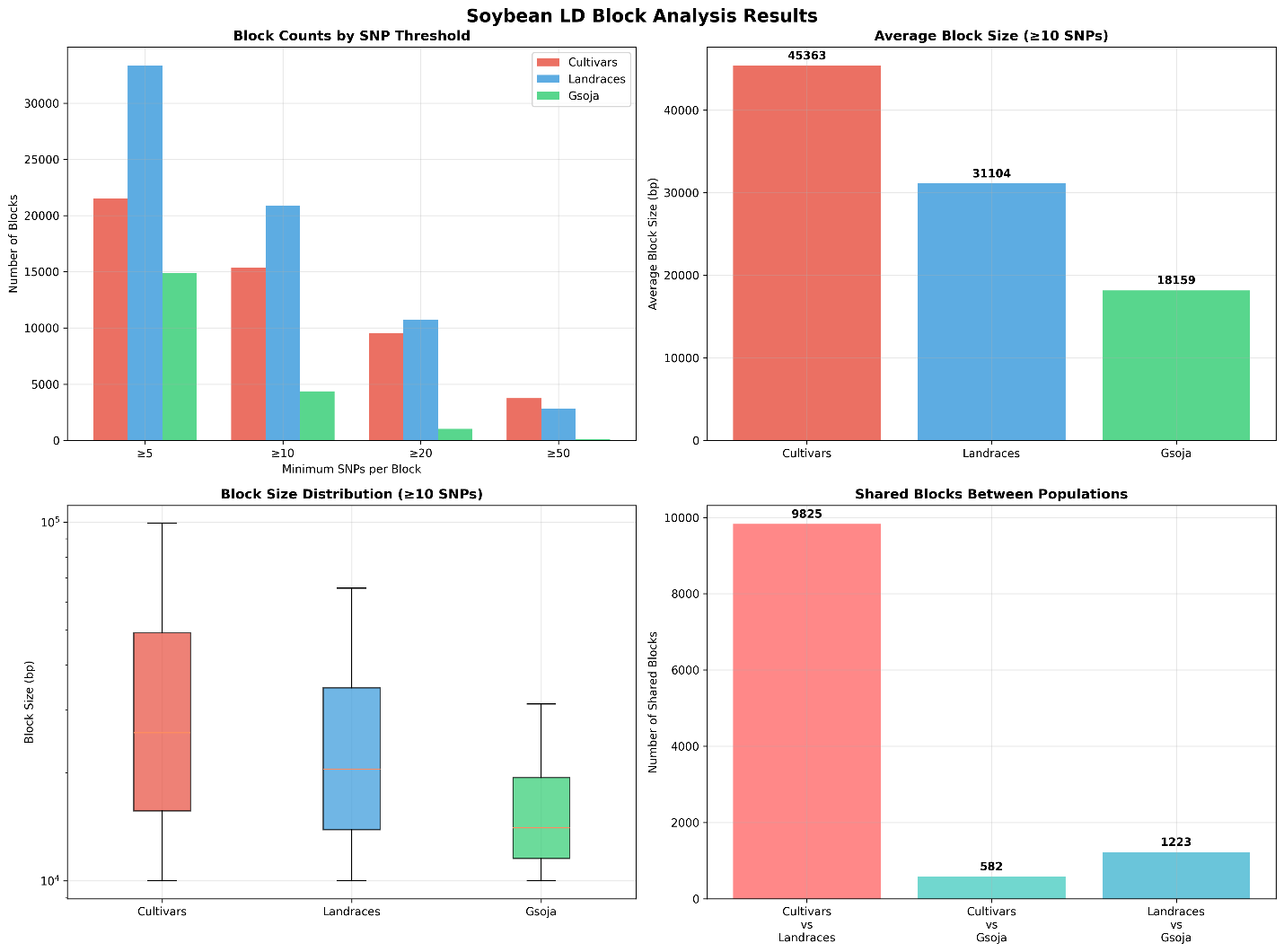

**Supplementary Figure S4: Count of Tagged SNPs per chromosome**

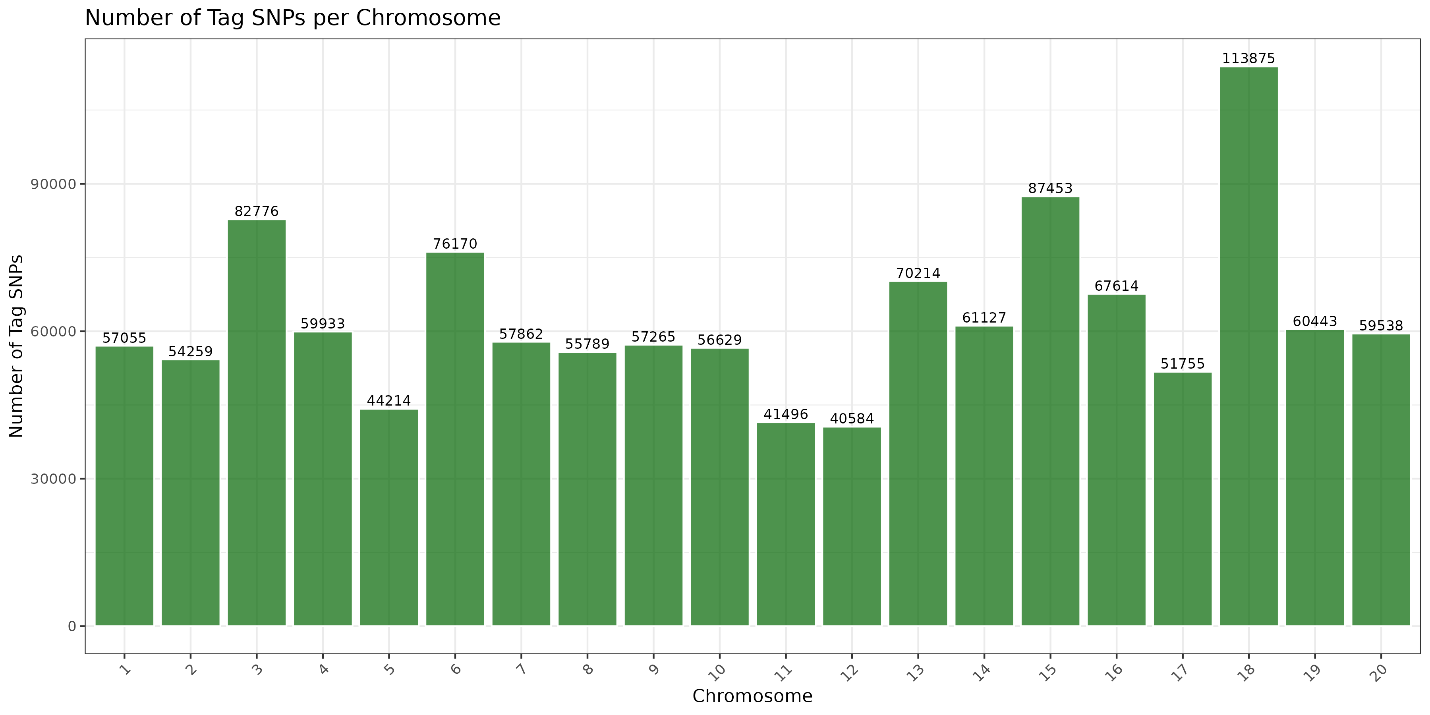

**Supplementary Figure S5: MAF distribution in the tagged SNPs set.**

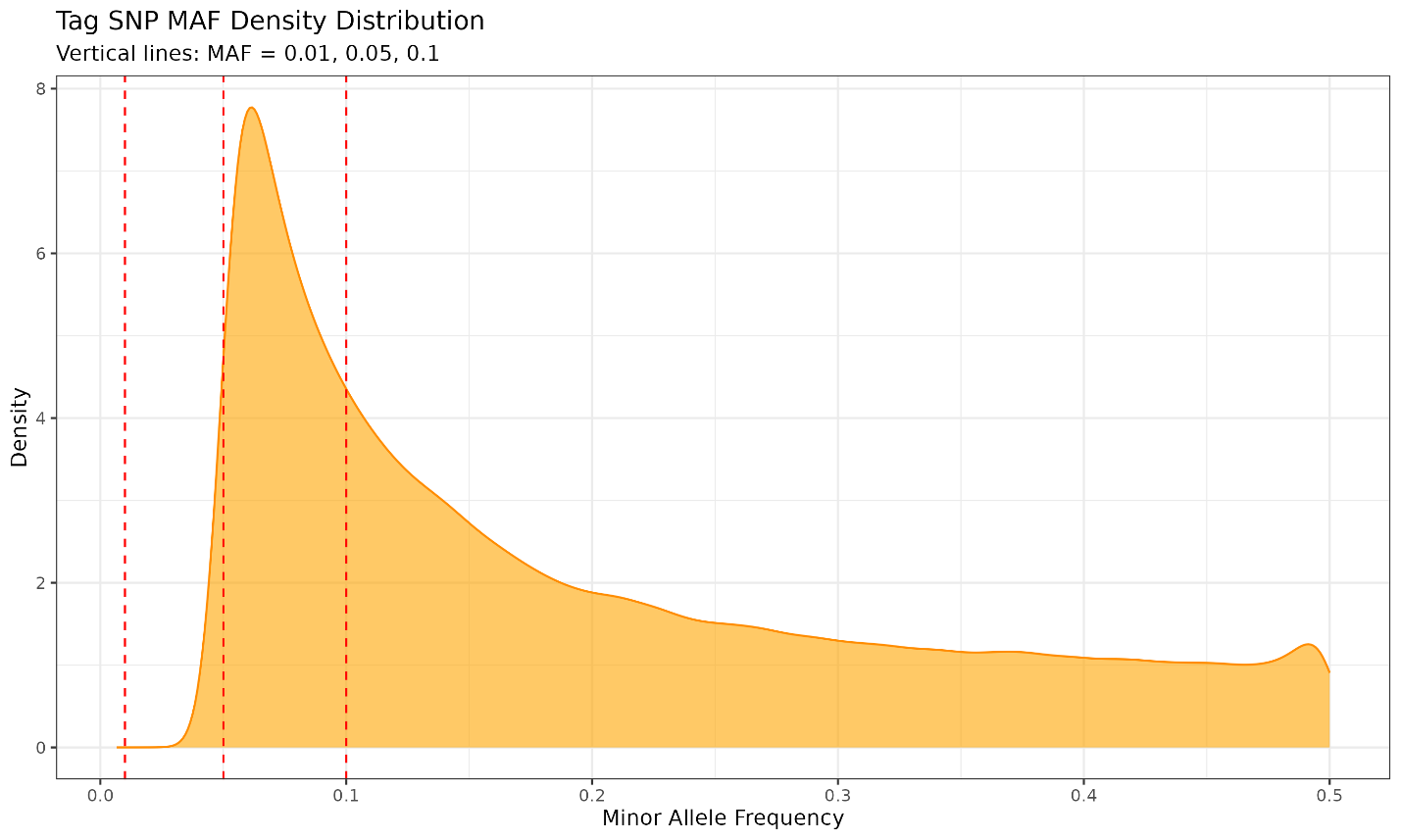

**Supplementary Figure S6: Manhattan plot for GWAS analysis of oil content**

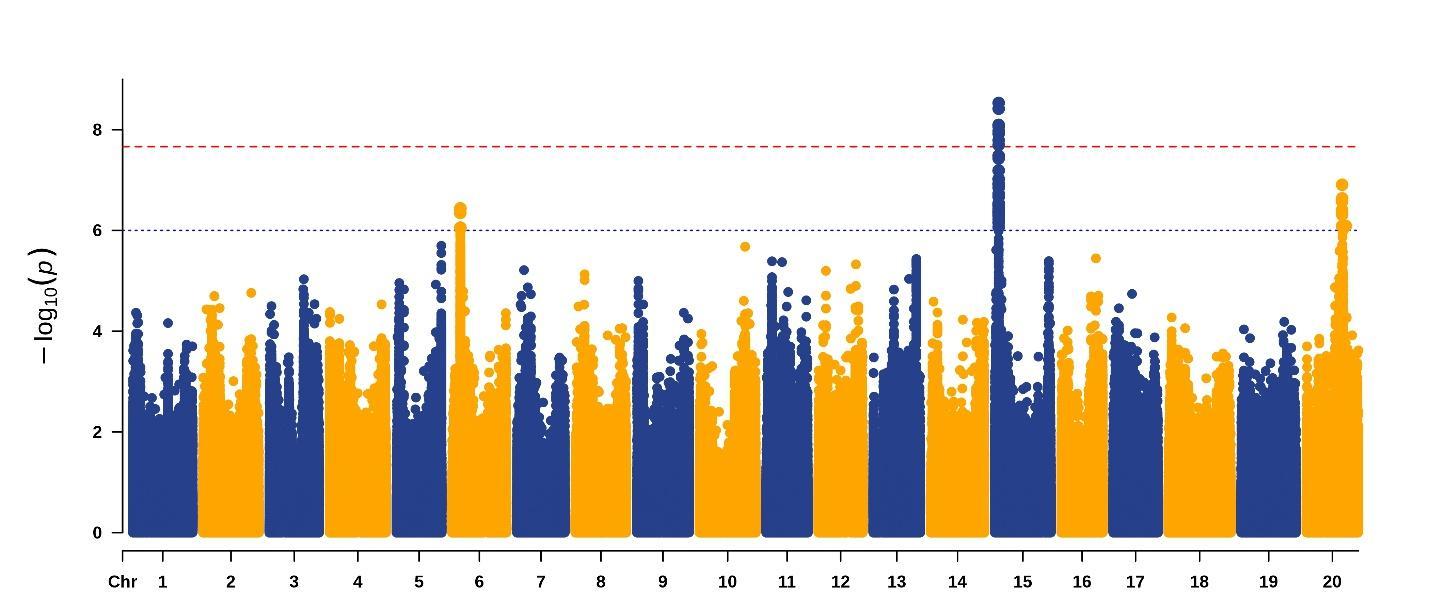

**Supplementary Figure S7: Manhattan plot for GWAS analysis of lodging**

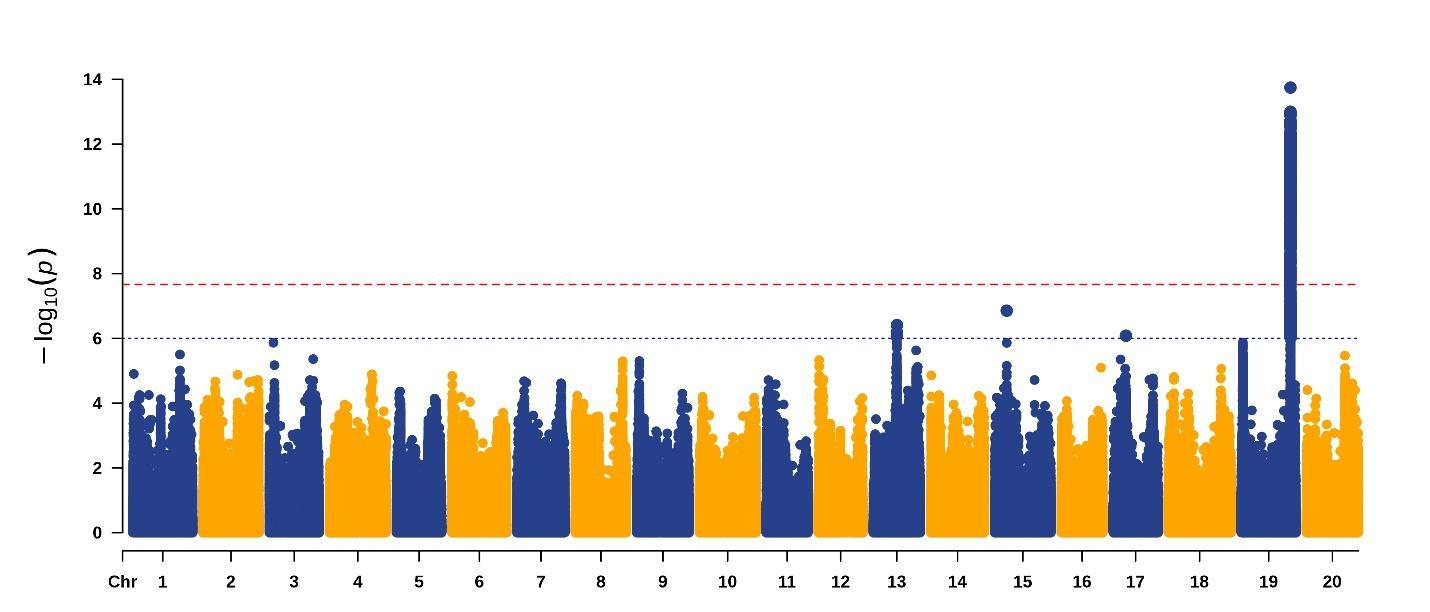

**Supplementary Figure S8: Manhattan plot for GWAS analysis of Palmitic acid**

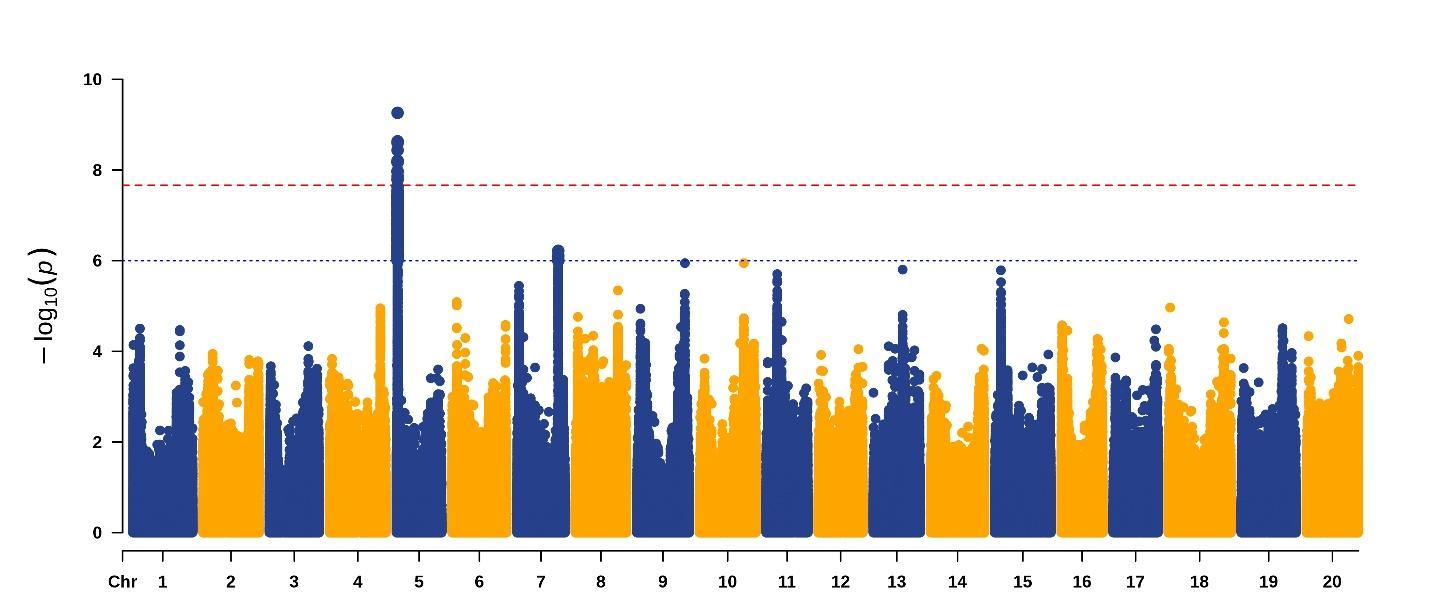

**Supplementary Figure S9: Manhattan plot for GWAS analysis of Stem growth habit**

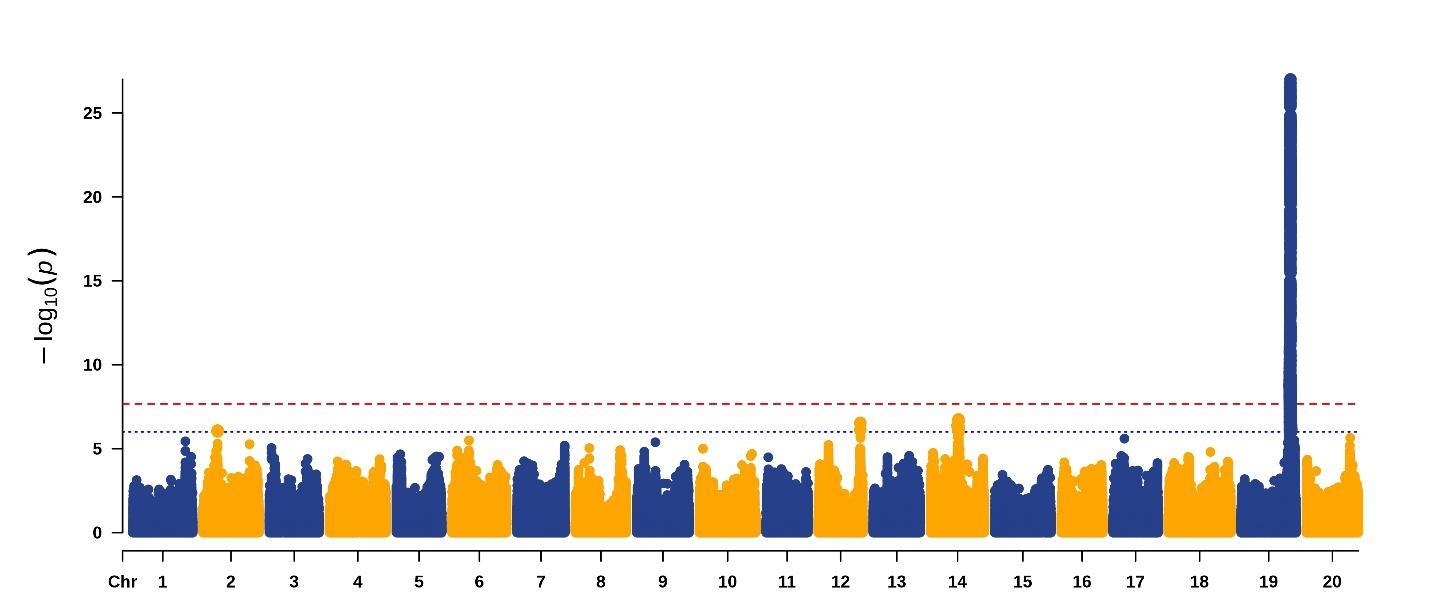

**Supplementary Figure S10: Haplo-pheno analysis of oil content QTL on Chr 15**

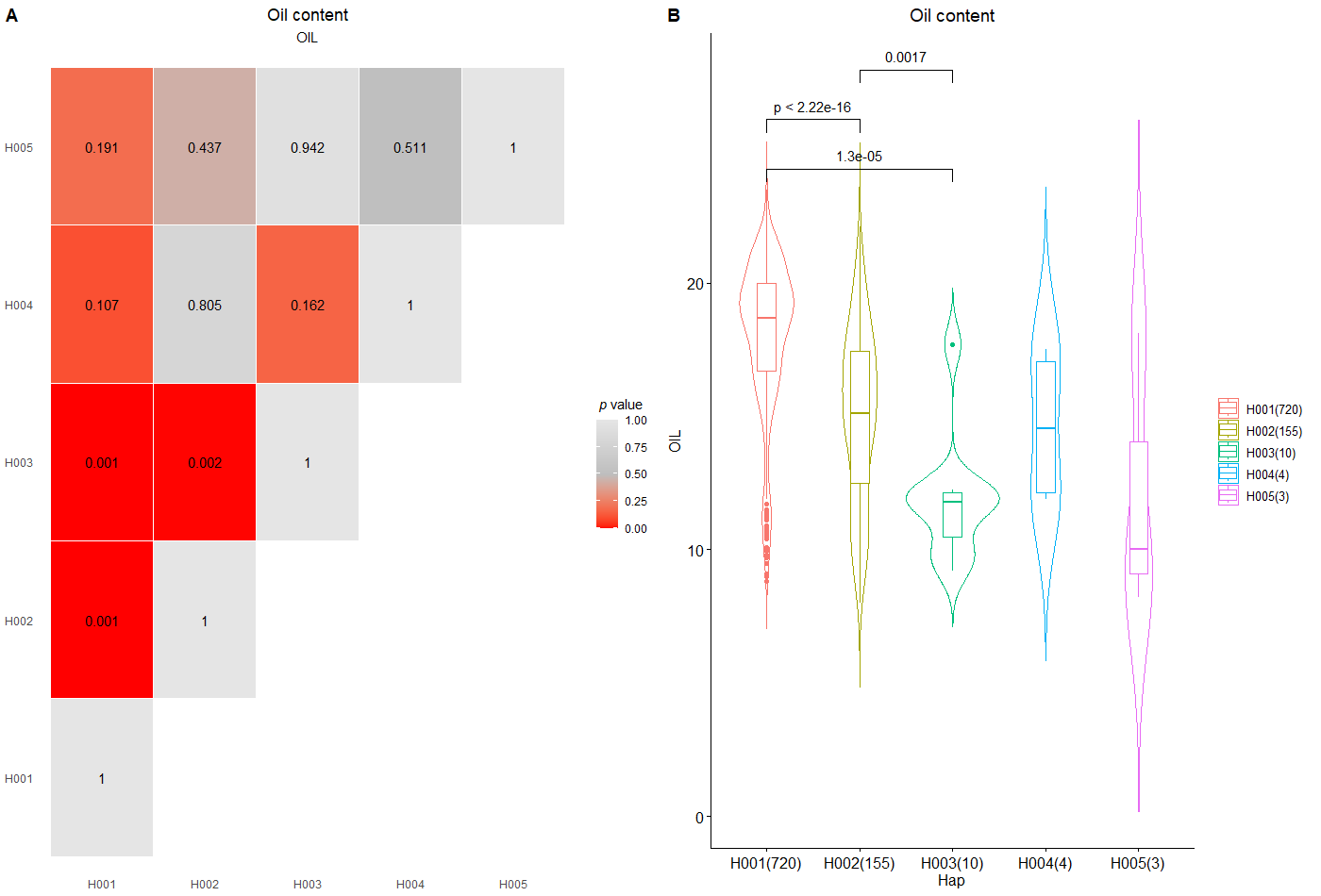

**Supplementary Figure S11: Distribution of major haplotypes for gene *Glyma.15G049200* (*Gm_41962*) across different countries**

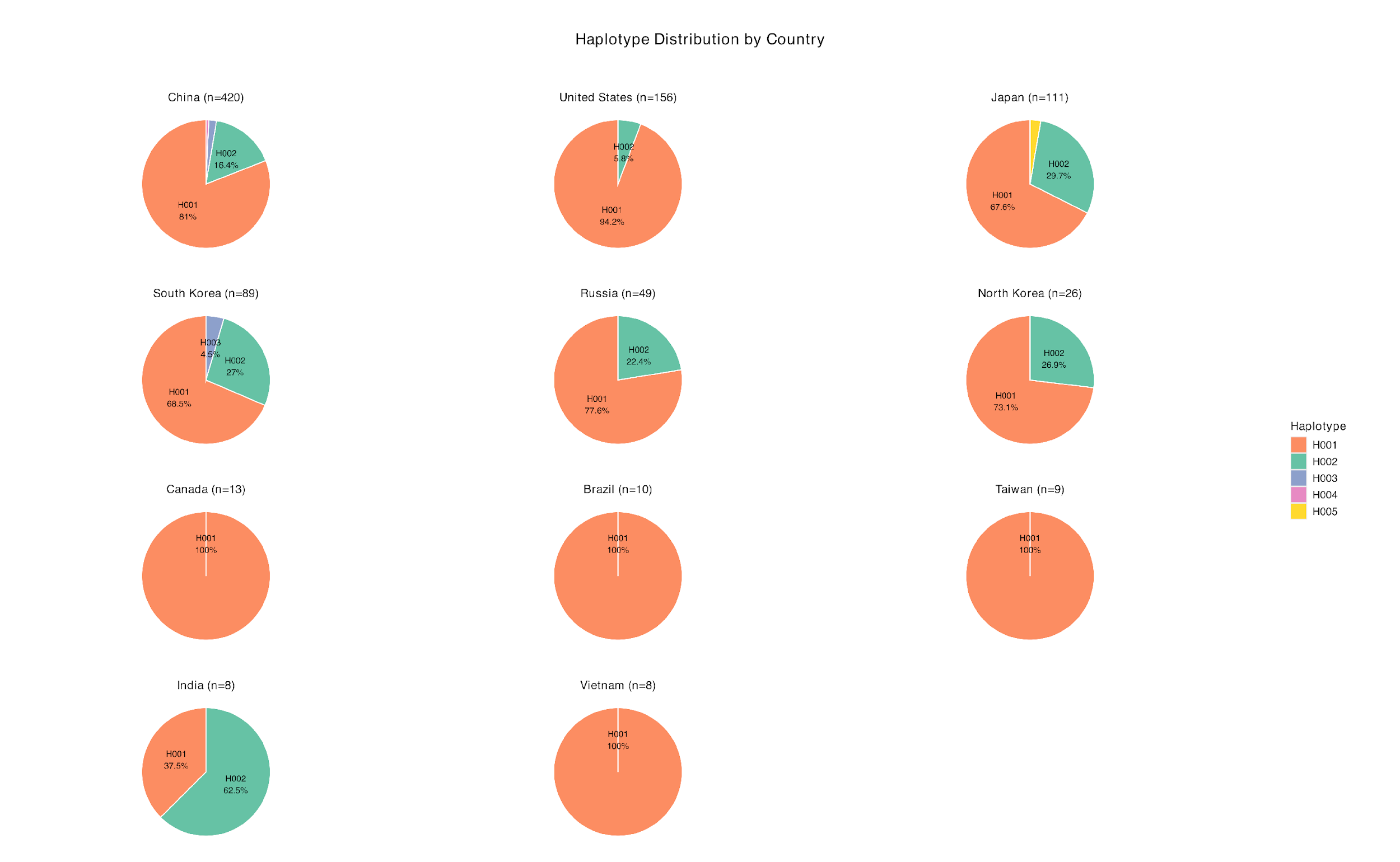

**Supplementary Figure S12: Association of haplotypes with SCN HG type 0 (race 3) phenotype**

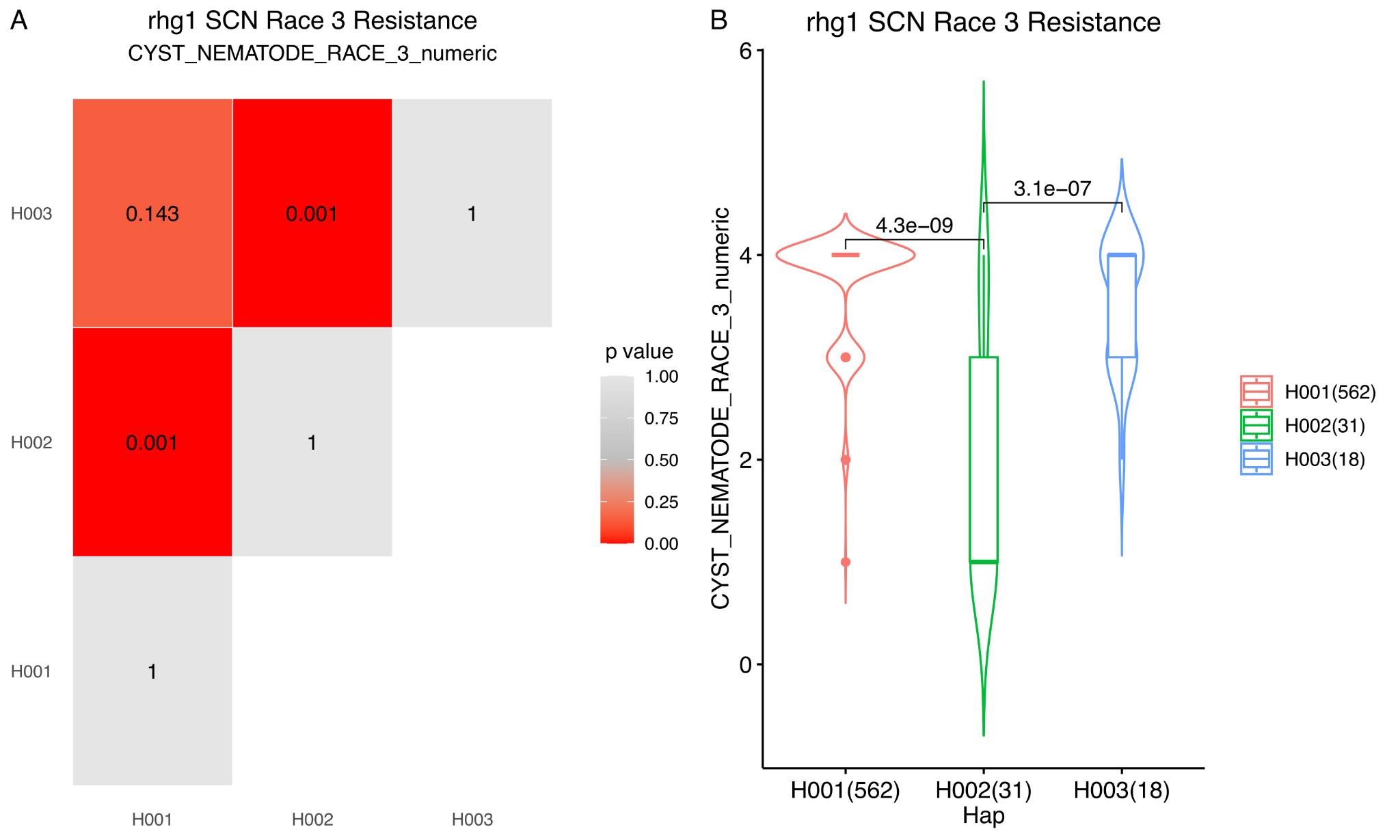

**Supplementary Figure S13: Country-wise haplotype distribution of the *rhg-1* gene**

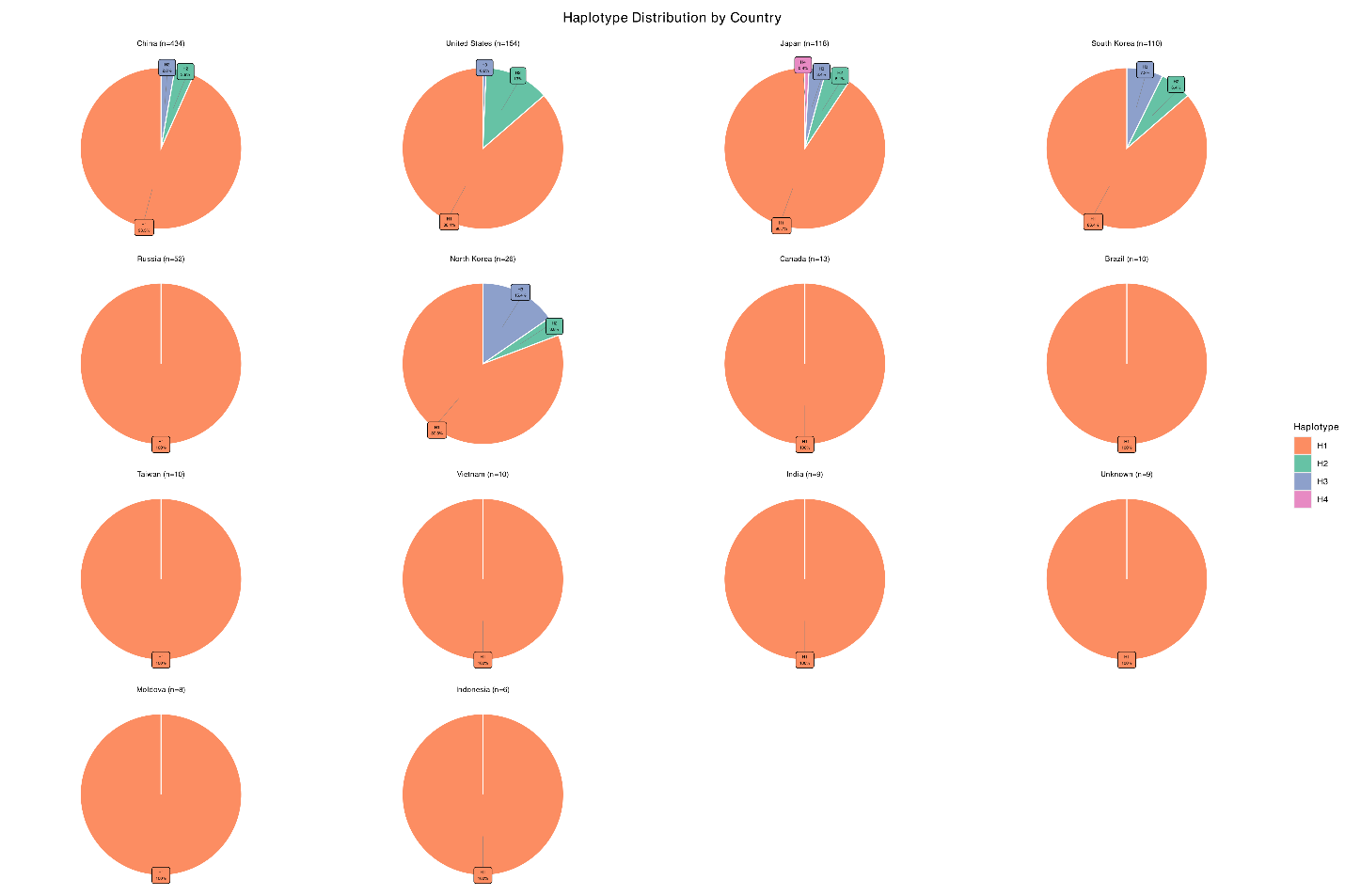

**Supplementary Figure S14:** Alluvial diagram illustrating the combinatorial effects of *rhg-1* and Rhg4 haplotypes and copy number variations on SCN race 5 resistance.

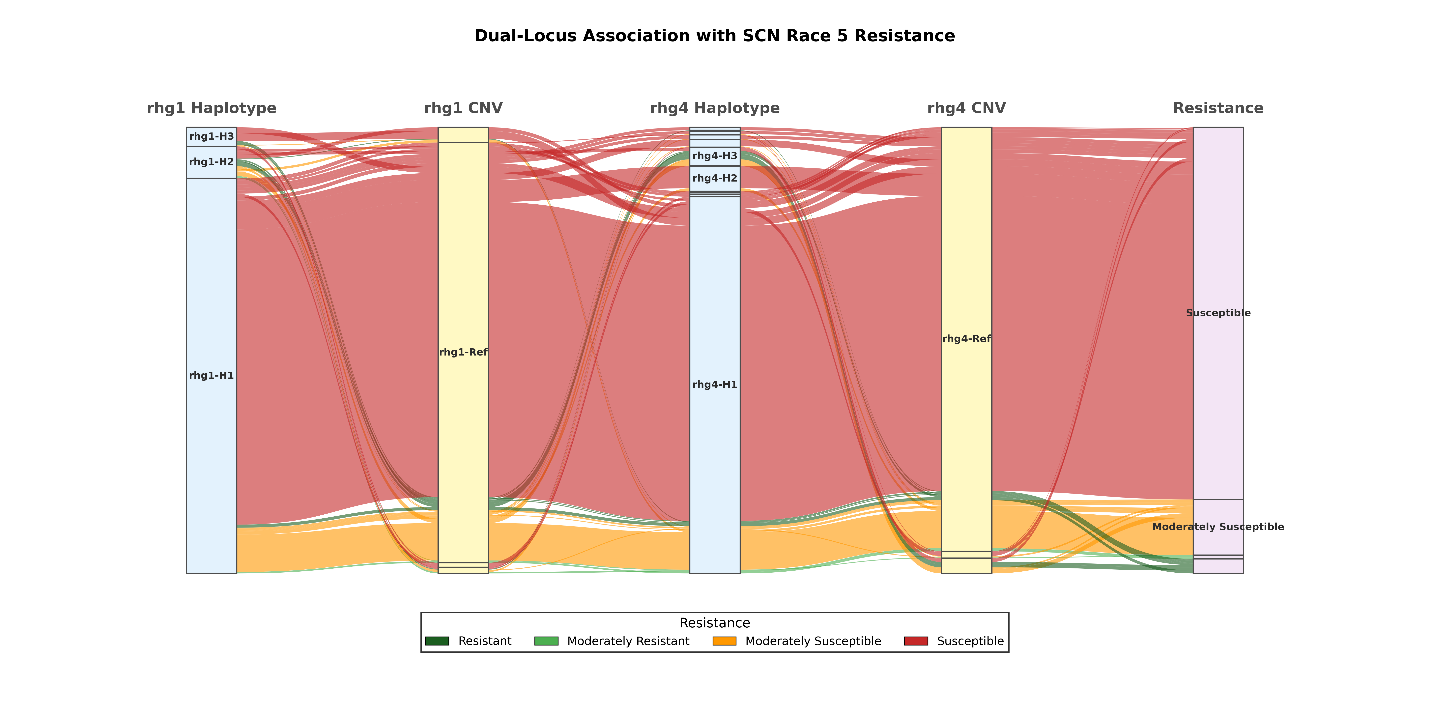

**Supplementary Tables**

**Supplementary Table S1:** Sequenced lines details with data estimate and metadata including species, origin and maturity group

**Supplementary Table S2:** Summary of mapping of resequencing data of 1278 soybean lines to the Wm82.a5 genome

**Supplementary Table S3:** Count of SNPs per chromosome without population-wide filters

| **Chromosome** | **No of SNPs** | **No of InDels** |
| --- | --- | --- |
| Chr01 | 3,106,233 | 307,852 |
| Chr02 | 2,491,528 | 279,877 |
| Chr03 | 2,659,889 | 290,806 |
| Chr04 | 2,672,083 | 271,850 |
| Chr05 | 2,230,703 | 239,395 |
| Chr06 | 2,813,262 | 315,772 |
| Chr07 | 2,471,485 | 281,567 |
| Chr08 | 2,221,100 | 288,760 |
| Chr09 | 2,573,117 | 278,614 |
| Chr10 | 2,686,877 | 294,899 |
| Chr11 | 1,885,079 | 224,260 |
| Chr12 | 2,089,257 | 225,338 |
| Chr13 | 1,972,553 | 291,464 |
| Chr14 | 2,921,351 | 292,195 |
| Chr15 | 3,090,639 | 329,871 |
| Chr16 | 2,205,709 | 254,498 |
| Chr17 | 2,124,458 | 249,578 |
| Chr18 | 3,493,626 | 379,772 |
| Chr19 | 2,602,261 | 274,582 |
| Chr20 | 2,667,568 | 268,726 |
| **Total** | **50,978,778** | **5,639,676** |

**Supplementary Table S4:** Metrics of number of SNPs identified in the soybean germplasm lines

**Supplementary Table S5:** Summary of genome-wide variant effects with MAF>0.05

| **Variant effect** | **Subcategory** | **Count of effect** | **Proportion of total SNPs (%)** |
| --- | --- | --- | --- |
| **Intergenic** |  | 9,600,721 | 84.47% |
| **Intronic** |  | 1,346,700 | 11.85% |
| **Splice site** |  | 27,254 | 0.24% |
| **Exonic** |  | 391,728 | 3.45% |
|  | Synonymous | 150,278 | 1.32% |
|  | Missense | 231,389 | 2.04% |
|  | Nonsense | 9,219 | 0.08% |
|  | Start/Stop disruption | 842 | 0.01% |
|  | Frameshift | 0 | 0.00% |
| **Other** |  | 4 | 0.00% |
| **Total** |  | **11,366,407** | **100.00%** |

**Supplementary Table S6:** Count for LOF SNPs on different soybean chromosomes

| **Chromosome** | **No of HIGH effect SNPs** | **Count fo LOF SNPs** |
| --- | --- | --- |
| Chr01 | 722 | 433 |
| Chr02 | 592 | 394 |
| Chr03 | 904 | 566 |
| Chr04 | 576 | 388 |
| Chr05 | 501 | 337 |
| Chr06 | 894 | 601 |
| Chr07 | 608 | 403 |
| Chr08 | 650 | 440 |
| Chr09 | 694 | 462 |
| Chr10 | 649 | 429 |
| Chr11 | 539 | 354 |
| Chr12 | 538 | 351 |
| Chr13 | 649 | 436 |
| Chr14 | 757 | 482 |
| Chr15 | 941 | 587 |
| Chr16 | 635 | 407 |
| Chr17 | 531 | 334 |
| Chr18 | 1165 | 728 |
| Chr19 | 636 | 400 |
| Chr20 | 660 | 418 |
| **Total** | **13,841** | **8,950** |

**Supplementary Table S7:** Phenotypic data on selected soybean lines for seed protein content, oil content and other traits obtained from the GRIN database **Supplementary Table S8:** MLM signals for protein content for GWAS on soybean individuals with rMVP (p-value ≤ 1 x 10-8) with SNP positions on Wm82.a5 reference genome **Supplementary Table S9:** MLM signals for oil content for GWAS on soybean individuals with rMVP (p-value ≤ 1 x 10-8) with SNP positions on Wm82.a5 reference genome **Supplementary Table S10:** MLM signals for lodging for GWAS on soybean individuals with rMVP (p-value ≤ 1 x 10-8) with SNP positions on Wm82.a5 reference genome **Supplementary Table S11:** MLM signals for palmitic acid content for GWAS on soybean individuals with rMVP (p-value ≤ 1 x 10-8) with SNP positions on Wm82.a5 reference genome
 **Supplementary Table S12**: MLM signals for stem growth habit for GWAS on soybean individuals with rMVP (p-value ≤ 1 x 10-8) with SNP positions on Wm82.a5 reference genome
 **Supplementary Table S13:** Haplotype sequence for each individual in the protein content associated gene *Glyma.15G049200* on Chr 15 **Supplementary Table S14:** Summary of phenotypic data range for protein content associated with each haplotype sequence **Supplementary Table S15:** Haplotype mapping of each individual in soybean germplasm collection across rhg1 and Rhg4 loci associated with SCN resistance **Supplementary Table S16:** Summary of haplotypes and CNV across soybean lines in both rhg1 and Rhg4 loci associated with SCN resistance in several races
 **Supplementary Table S17:** Haplotype and CNV combinations with expected phenotypes

**Supplementary Table S18:** Training set for genomic prediction of multi class variations for rhg1 locus

**Supplementary Table S19:** Test set for genomic prediction of multi class variations for rhg1 locus
